# SubCellSeg: A generalist model for subcellular structure segmentation in 3D fluorescence microscopy

**DOI:** 10.64898/2026.11.10.765799

**Authors:** Suraj Mishra, Ashwin Samudre, Amil Khan, Antoine Borensztejn, Caroline Hookway, Derek J. Thirstrup, Jianxu Chen, Susanne M. Rafelski, Matheus P. Viana

## Abstract

Three-dimensional (3D) segmentation of subcellular structures from fluorescence microscopy images is the foundation of many analyses in quantitative cell biology. The prevailing approach addresses 3D segmentation one structure at a time, with a separate model trained for each structure type. By developing SubCellSeg, we demonstrate that a single model, trained jointly across a diverse range of structures, substantially and consistently outperforms structure-specific specialist models. SubCellSeg is a 3D segmentation model trained on expert-curated fluorescence images and their computationally generated *reference masks*, of 13 morphologically distinct subcellular structures in human induced pluripotent stem (hiPS) cell lines from the Allen Cell Collection, spanning compact puncta, planar junctions, reticular networks, and tubular structures. Without any structure-specific input, SubCellSeg achieves a mean Dice score of 0.909 across the 13 structures and outperforms each of the 13 specialist models on the structure it was trained for (specialist mean Dice 0.773). SubCellSeg also achieves zero-shot generalization, segmenting subcellular structures with morphologies not represented in the training data. This generalization extends to images acquired on different microscopes and with different imaging configurations, to cells under drug perturbation, and to a morphologically distinct cell type, hiPS cell-derived cardiomyocytes. Where zero-shot results are insufficient for a specific application, SubCellSeg can be fine-tuned: starting from its pretrained weights, 10% of the labeled images (16 images) gave better results than training a new model from scratch on all of them. SubCellSeg is distributed through the napari plugin `allencell-segmenter-ml` [1] with pretrained checkpoints, so it can be run on new data without any training; the training and evaluation code is available at https://github.com/AllenCell/SubCellSeg.

## Introduction

Most quantitative analyses of subcellular organization begin by deciding which voxels in an image belong to a given structure. This segmentation step enables measurements of structure morphology, quantification of structural responses to perturbation, and construction of integrative models of cellular organization [2, 3]. The extreme diversity of subcellular structures makes the problem challenging, e.g., mitochondria form interconnected tubular networks [4, 5]; gap junctions cluster into small plaques at cell-cell contacts [6]; nuclear pore complexes appear as roughly uniform puncta on the nuclear envelope [7]; the endoplasmic reticulum combines flat sheets, concentrated near the nucleus, with a highly dynamic network of tubules in the cell periphery, where densely packed tubules can appear as sheets at conventional optical resolution [8]. These structures differ substantially in their size and morphology and, therefore, once imaged via fluorescently tagged proteins, vary in their image properties used for segmentation, including signal statistics, contrast, density, scale, and boundary definition. These differences are amplified in 3D, where resolution along the optical axis is several-fold worse than in the image plane and out-of-focus light blurs structure boundaries.

Segmentation of subcellular structures long predates deep learning, and the classical methods established a convention that deep learning-based methods later inherited: one method per structure. Many of the first methods were designed for 2D images and later extended to 3D stacks. Early approaches relied on intensity thresholding and morphological cleanup [9], which works when a structure is bright, compact, and well separated. Structures that are none of those things led to filters tuned to particular shapes: wavelet multiscale products for punctate structures such as vesicles and endosomes [10], and Hessian ridge filters, borrowed from vascular imaging, for tubular and reticular ones such as mitochondria and the ER [11, 12]. Combining these filters with structure-specific pre- and post-processing produced dedicated tools for single structures, such as MitoGraph, which reconstructs the mitochondrial network in 3D [13], and CiliaQ, which quantifies ciliary morphology and fluorescence in 2D, 3D and 4D images [14]. Each performs well on the structure it was built for and transfers to no other. Methods aimed more broadly at subcellular structures followed, coupling deconvolution to a convex optimization-based segmentation of the structures themselves [15], while open platforms turned these steps into reproducible 3D structure workflows [16, 17]. These were advances in reproducibility rather than in generality: Squassh assumes locally homogeneous intensities and several parameters to be set per image [15]; TANGO targets nuclear compartments specifically [17]. Output remained hyperparameter-dependent and tuned per structure and often per dataset. Interactive pixel classifiers trained on sparse user annotation offer a usable mask for a fraction of the effort of dense manual labeling [18], but the classifier is still trained anew for each structure.

The Allen Cell and Structure Segmenter [1] is the most extensive collection of such structure-specific workflows for 3D intracellular segmentation: curated classical image-processing workflows, and deep learning workflows trained on their outputs, for over 20 intracellular structures. Deep learning has largely retained this per-structure organization, with a separate network usually trained for each structure or task of interest [19]. Whether classical or deep-learning-based, these methods share one organizing principle: each structure receives its own model or workflow. This works well when a single structure is the focus of a study, but it does not scale. Developing one model per structure duplicates training pipelines, hyperparameter choices, annotation efforts, and failure modes, and keeping each model working as imaging conditions change repeats that cost. It also prevents models from sharing information across structures. Images of different structures in the same cell type share the same cellular context, and when acquired under a single imaging protocol, they may also share many fluorescence-image features, such as foreground-background contrast, noise statistics, voxel size, and anisotropic resolution. A model trained across multiple structures may therefore learn reusable representations that transfer across subcellular targets.

*Generalist* deep learning segmentation models comprising single models with one set of weights that segment many different kinds of images, have already proven useful for whole-cell and nuclear segmentation, as illustrated by Cellpose [20, 21] and Mesmer [22], and related ideas underlie recent foundation models for biological imaging [23–25]. However, 3D subcellular fluorescence segmentation differs substantially from the whole-cell or nuclear segmentation targeted by these other approaches: the targets are smaller, more morphologically diverse, often sparse or reticular, and embedded in anisotropic volumetric images. Here, we introduce SubCellSeg, a single 3D model trained jointly on images of 13 endogenously tagged subcellular structures from cell lines in the Allen Cell Collection [2, 26] spanning very diverse morphologies, and resulting in a generalist, morphology-agnostic segmentation model: one set of weights for all structures, in contrast to the conventional *specialist* approach of one model per structure. SubCellSeg receives no structure label at training or inference time, and is trained to segment any structure that produces a fluorescence contrast pattern resembling those seen during training.

To evaluate SubCellSeg, we compare a single jointly trained model against structure-specific specialist models trained under an identical protocol. We also benchmark SubCellSeg against established segmentation tools: general-purpose methods trained on the same data or used off the shelf, published tools built for individual structure classes (ER networks, cell boundaries, diffraction-limited spots), and a promptable 3D foundation model. Beyond that comparison, we ask three practical questions: 1)Does the segmentation quality of the jointly trained model transfer to images acquired on different microscopes, at different magnifications, or in different cell types? 2) Is the segmentation quality the same when cells undergo perturbations, and the structure shape deviates substantially from the training distribution? 3) When images of a structure with a morphology not seen in training are acquired,how many labeled volumes are needed for fine-tuning? Our results suggest that a multi-structure training strategy yields scalable, generalist behavior for 3D subcellular segmentation: SubCellSeg can be used as is for a first segmentation pass, and then further fine-tuned on a small labeled set when that is not sufficient.

## Results

### A single SubCellSeg model segments 13 diverse subcellular structures

SubCellSeg was trained on a strategically selected subset of the WTC-11 hiPSC Single-Cell Image Dataset v1 [2] (hereafter, the WTC-11 dataset), a publicly available resource of WTC-11 hiPS cells imaged at high resolution on a spinning-disk confocal microscope (SDCM-1; Methods). Each cell line used for training the model carries an endogenously tagged mEGFP fused to a protein marking a subcellular structure of interest, avoiding the morphological artifacts of overexpression.

The WTC-11 dataset contains raw microscopy images and corresponding reference masks for 25 different subcellular structures. Reference masks were generated using the classic image segmentation workflows in the Allen Cell and Structure Segmenter. Biologists reviewed and assessed the quality of these reference masks on a per-structure basis [2]. Conceptually, the entire dataset of paired fluorescence and binary segmentation images could be used for training SubCellSeg. However, strategically, twelve structures were excluded from the training set due to : (a) systematic suboptimal segmentation for certain structure segmentation workflows that would introduce noise into the training set, (b) structures with context-dependent morphology that limits generalizability, and (c) ambiguous labels arising from multiple segmentation targets for the same fluorescent structure. The remaining 13 structures spanned a wide spectrum of morphological categories needed to create a generalist model and were judged to have sufficiently accurate reference masks as reliable training annotations. The 13 structures used for training were actin bundles, actomyosin bundles, adherens junctions, desmosomes, gap junctions, tight junctions, the ER (two markers), the Golgi, endosomes, mitochondria, nuclear pores, and nuclear speckles (Fig. 1a). Training on these paired image and reference masks avoids the laborious and expensive step of manual 3D annotation. For each structure, 267 z-stacks (FOVs; the minimum available across the 25 structures in the WTC-11 dataset) were split into non-overlapping sets for model training (160 samples), validation (40 samples), and testing (67 samples, curated to 57; see Methods and Supplementary Note 1), ensuring an equal number of images per structure.

**Figure 1:**
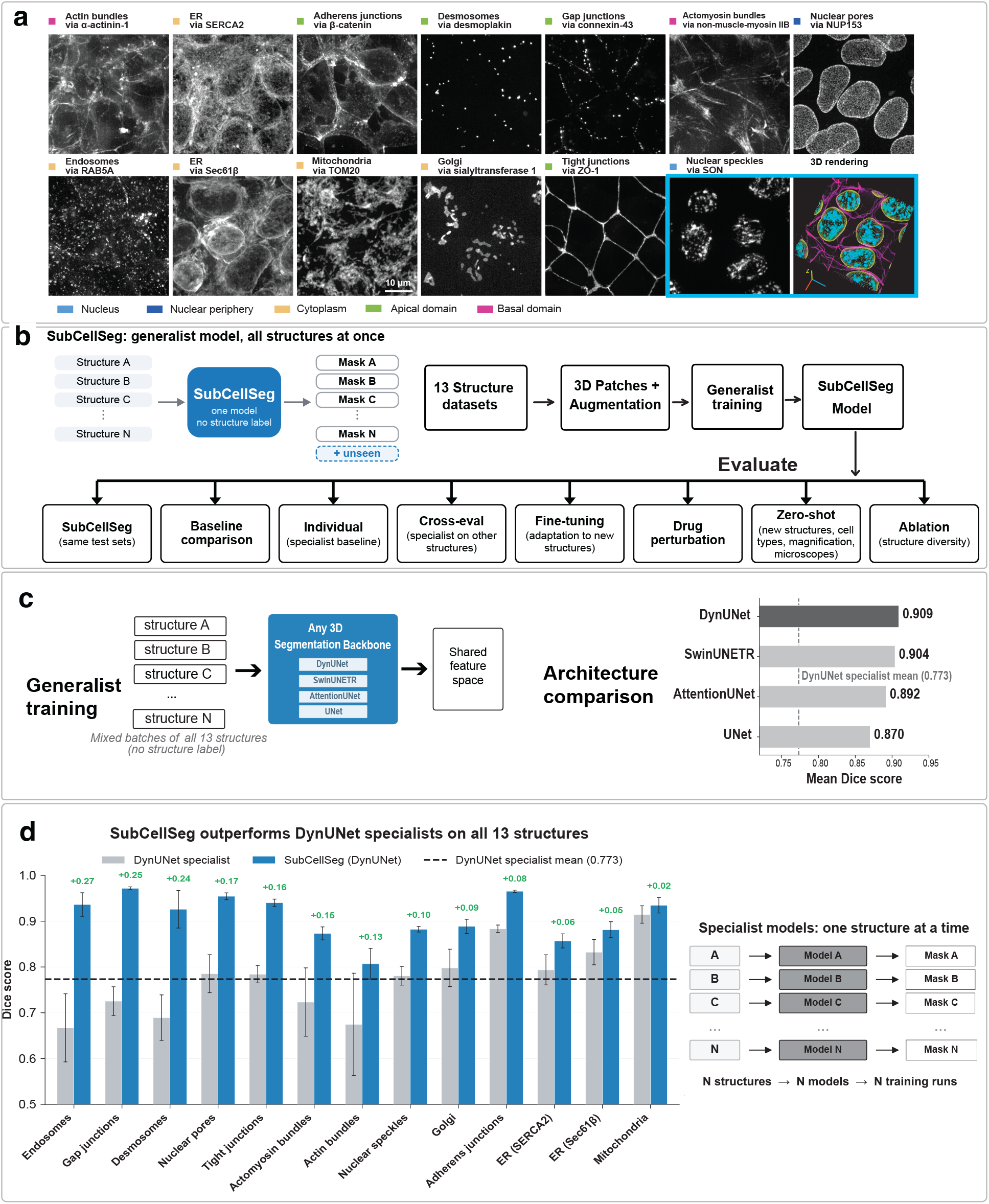
**SubCellSeg: a generalist model for 3D segmentation across 13 subcellular structures**. (a) Maximum-intensity projections of the 13 structures used to train SubCellSeg, labeled as structures via tagged protein and color-coded by subcellular location (nucleus, nuclear periphery, cytoplasm, apical domain, basal domain). Each cell line carries an endogenously tagged mEGFP fusion to the indicated protein in the WTC-11 hiPSC background. Boxed, right: VolE 3D rendering of the nuclear speckle z-stack (yellow) together with the nuclear (cyan) and cell membrane (magenta) channels for context, illustrating the volumetric nature of the data. (b) Left, the SubCellSeg paradigm: a single model receives images of all structures, with no structure label, and returns a mask for each, including structures not seen during training. Right, evaluation workflow: images from the 13 structure image datasets are cropped into augmented 3D patches for generalist training, and the resulting model is evaluated on the held-out test sets, against existing segmentation methods and individual DynUNet specialists, by cross-evaluation of specialists on other structures, by fine-tuning to new structures, under drug perturbation, by zero-shot application to new structures, cell types, magnifications and microscopes, and by ablation of training-set structure diversity. (c) Left, generalist training: mixed batches containing all 13 structures are fed to a single 3D segmentation backbone (DynUNet, SwinUNETR, AttentionUNet or UNet), yielding a shared feature space. Right, architecture comparison: mean Dice across the 13 structures for each backbone, compared with the DynUNet specialist mean (dashed line, 0.773); the *x*-axis starts at 0.72. DynUNet (0.909) is the default backbone. (d) Left, per-structure Dice for the DynUNet specialists (grey) and SubCellSeg with a DynUNet backbone (blue) on the held-out test set. Error bars: standard deviation across test images. Green numbers above the bars give the absolute Dice gain of SubCellSeg over the specialist for each structure; SubCellSeg outperforms the specialist on all 13 structures (mean gain +0.136). Right, the conventional specialist paradigm, in which N structures require N separately trained models.

SubCellSeg interleaves images from all 13 structures into mixed batches and feeds them to a single 3D segmentation backbone in a structure-agnostic way (Fig. 1b, left). The model outputs a single-channel probability map; it is never provided with information on which structure it is looking at and has no structure-specific output. To test the effectiveness of this design, we evaluated SubCellSeg in a series of experiments (Fig. 1b, right), starting with the choice of backbone. We trained four backbones in the same way (Fig. 1c): DynUNet [27], SwinUNETR [28], AttentionUNet [29] and a standard 3D UNet [30] (details in Methods).

All four backbones reached high accuracy when trained on the 13 structures (Fig. 1c): DynUNet (0.909) *>* SwinUNETR (0.904) *>* AttentionUNet (0.892) *>* standard UNet (0.870). The top three lay within 0.02 Dice of one another, so with multi-structure training the choice among these backbones mattered little. The DynUNet’s slight edge and lower memory footprint relative to SwinUNETR made it our default architecture for SubCellSeg, and all results below use it unless stated otherwise. The standard UNet’s *∼*4 percentage-point lag behind DynUNet (0.870 vs. 0.909; Supplementary Table 3) is consistent with the benefit of dynamic patch-resampling, anisotropic kernels, and attention mechanisms when input scales vary across structures. The per-structure breakdown of all four backbones is provided in Fig. 7. Backbone choice mattered most for ER via SERCA2, ER via Sec61*β* and nuclear pores via NUP153, where the standard UNet trailed the other backbones by up to 0.11 Dice, and least for Golgi via sialyltransferase 1 and desmosomes via desmoplakin, where all four backbones lay within 0.011 Dice of one another.

### SubCellSeg outperforms structure-specific specialist models

To test whether a single generalist model trained on 13 structures outperforms the conventional specialist pipeline in which each structure receives its own dataset, model, and training run (Fig. 1d, right), we trained 13 structure-specific DynUNet specialist models (DynUNet only, the default SubCellSeg backbone; Methods) with identical hyperparameters and number of epochs, then evaluated both on the same held-out test sets. The specialist 13-DynUNet model ensemble reached a mean Dice score (overlap between predicted and reference masks, from 0 to 1; Methods) of 0.773 *±* 0.06 while SubCellSeg with a DynUNet backbone reached 0.909, an absolute improvement of 0.136 Dice (13.6 percentage points; shown in Fig. 1c, d and Table 1). The improvement was consistent across different structures with SubCellSeg outperforming all 13 specialist models with a range of improvement of +0.02 Dice (mitochondria via TOM20, where the specialist model was already performing well) to +0.27 (endosomes via RAB5A, where segmenting small punctate signals is more challenging for the specialist model). Randomly selected test images for four structures are shown in Fig. 2 (the remaining nine structures are shown in Extended Data Fig. 1).

**Figure 2:**
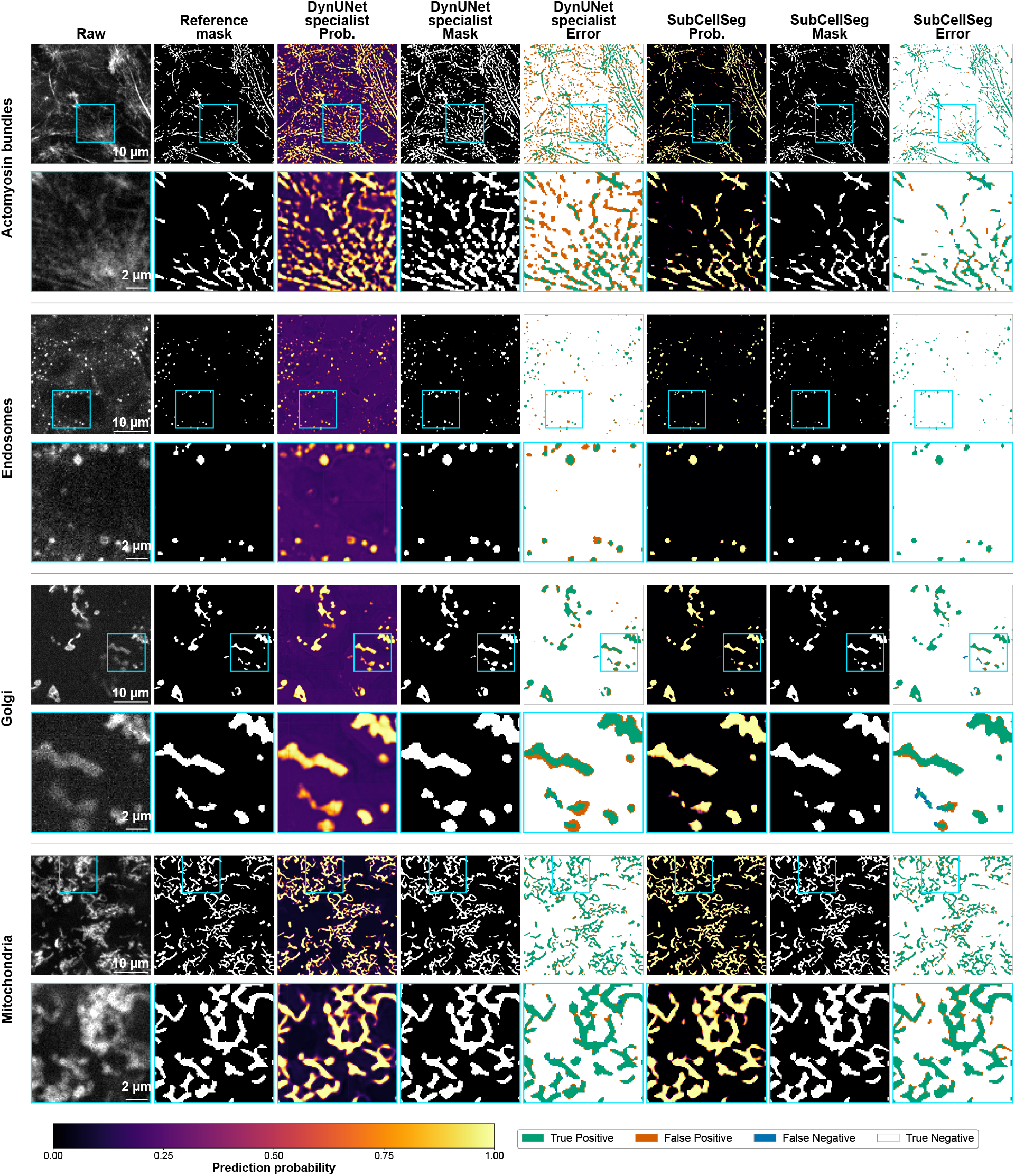
Qualitative segmentation results for four of the 13 trained structures. Random single *z*-slices from 3D image stacks for actomyosin bundles via non-muscle myosin IIB, endosomes via RAB5A, Golgi via sialyltransferase 1 and mitochondria via TOM20; the remaining nine structures are shown in Extended Data Fig. 1. Each structure occupies two rows: the full field of view (top) and a 3.2*×* magnification of the cyan boxed region (bottom), placed where the DynUNet specialist and SubCellSeg masks differ most. Columns show the raw image, the reference mask used as the training and evaluation target, the DynUNet specialist probability map, binary mask and error map, followed by the SubCellSeg probability map, binary mask and error map. Probability maps share the colour scale at bottom (prediction probability 0–1). Error maps are color-coded: green, true positive; orange, false positive; blue, false negative; white, true negative. Scale bars, 10 *µ*m (full view) and 2 *µ*m (magnified view).

**Table 1:**
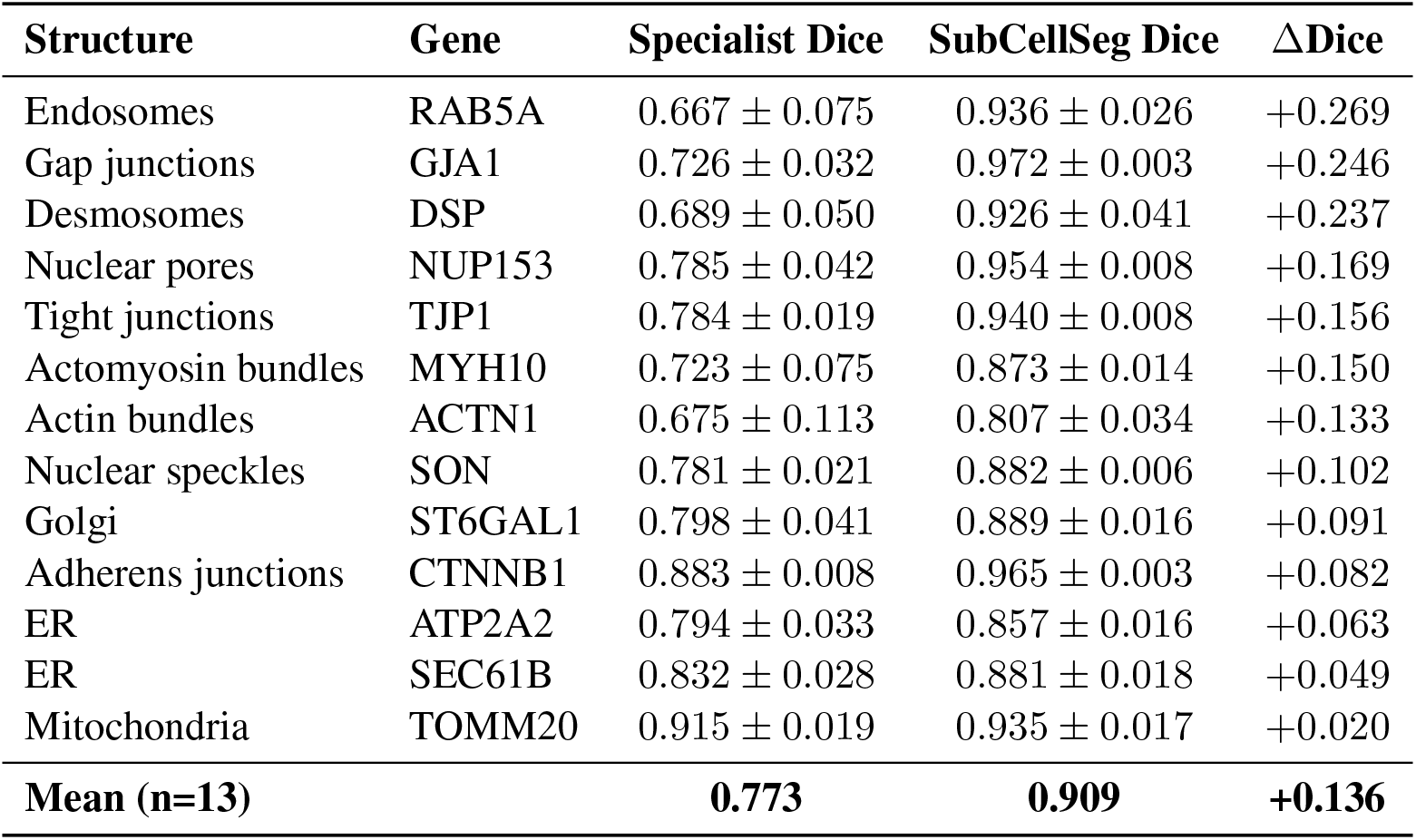
Per-structure segmentation performance. Mean Dice score *±* standard deviation over the 57 held-out test images per structure for the specialist DynUNet, SubCellSeg DynUNet (default), and the absolute Dice gain. All values are mean Dice over the test images for each structure. The mean over the 13 structures is reported in the bottom row.

| **Structure** | **Gene** | **Specialist Dice** | **SubCellSeg Dice** | **ΔDice** |
|---|---|---|---|---|
| Endosomes | RAB5A | 0.667 ± 0.075 | 0.936 ± 0.026 | +0.269 |
| Gap junctions | GJA1 | 0.726 ± 0.032 | 0.972 ± 0.003 | +0.246 |
| Desmosomes | DSP | 0.689 ± 0.050 | 0.926 ± 0.041 | +0.237 |
| Nuclear pores | NUP153 | 0.785 ± 0.042 | 0.954 ± 0.008 | +0.169 |
| Tight junctions | TJP1 | 0.784 ± 0.019 | 0.940 ± 0.008 | +0.156 |
| Actomyosin bundles | MYH10 | 0.723 ± 0.075 | 0.873 ± 0.014 | +0.150 |
| Actin bundles | ACTN1 | 0.675 ± 0.113 | 0.807 ± 0.034 | +0.133 |
| Nuclear speckles | SON | 0.781 ± 0.021 | 0.882 ± 0.006 | +0.102 |
| Golgi | ST6GAL1 | 0.798 ± 0.041 | 0.889 ± 0.016 | +0.091 |
| Adherens junctions | CTNNB1 | 0.883 ± 0.008 | 0.965 ± 0.003 | +0.082 |
| ER | ATP2A2 | 0.794 ± 0.033 | 0.857 ± 0.016 | +0.063 |
| ER | SEC61B | 0.832 ± 0.028 | 0.881 ± 0.018 | +0.049 |
| Mitochondria | TOMM20 | 0.915 ± 0.019 | 0.935 ± 0.017 | +0.020 |
| **Mean (n=13)** |  | **0.773** | **0.909** | **+0.136** |

SubCellSeg also predicted its segmentation masks with more confidence than the specialist models. Its probability maps were near-binary, with background driven close to zero and foreground close to one, whereas the specialist models assigned appreciable probability to background and produced blurry foreground-background transitions (examples in Fig. 2). As a result, SubCellSeg’s masks are largely insensitive to the exact binarization threshold. All binary masks reported here use a fixed threshold of 0.5 (Methods).

The gain was largest where the specialists were weakest (Fig. 1d; Table 1). The five structures with the lowest specialist Dice (endosomes, actin bundles, desmosomes, actomyosin bundles and gap junctions; 0.67–0.73) each gained at least 0.13 Dice and reached 0.81–0.97 with SubCellSeg. Structures that the specialists already segmented well gained less: adherens junctions (+0.08, from 0.88), the two ER markers (+0.05 and +0.06) and, smallest of all, mitochondria (+0.02, from 0.915). SubCellSeg therefore also narrowed the spread across structures, from 0.67–0.92 Dice for the specialists to 0.81–0.97. The error maps in Fig. 2 show where the two models differ.

If specialist models learn features specific to their own structure, they may perform poorly on images of other structures. To test this, we applied each of the 13 specialist DynUNet backbone models to all 13 test structures, yielding a 13 *×* 13 Dice matrix (heatmap shown in Fig. 3a). Specialists scored a mean Dice of 0.77 on their own structure but only 0.50 on the other twelve (range 0.02–0.82), with the steepest drops for sparse targets such as desmosomes and endosomes. Each specialist model was exposed to a single structure during training and the information it learned could therefore be structure-specific. SubCellSeg, by contrast, never sees a structure-specific label or identifier; its predictions depend on image content alone, and it achieves the highest Dice on every structure (Fig. 3b, top row).

**Figure 3:**
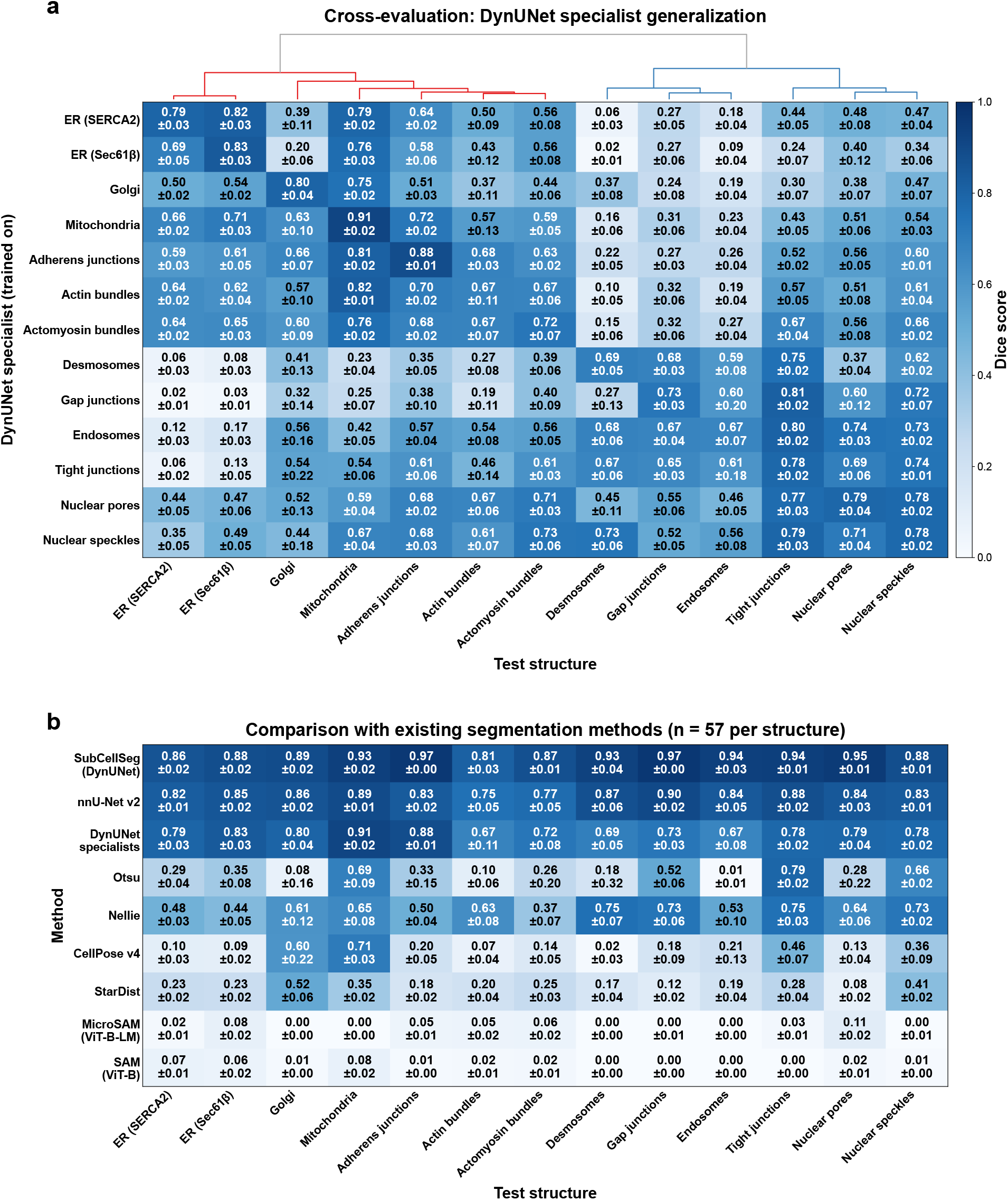
**Specialist models do not generalize across structures**. (a) Cross-evaluation matrix: Dice for each of 13 structure-specific specialist DynUNet models (rows) applied to each of 13 structures (columns), mean ± s.d. across the held-out test set. Diagonal entries are each specialist on its own target; off-diagonal entries show performance on structures it was not trained for. Hierarchical clustering on rows and columns separates the structures into two groups. (b) SOTA method comparison: per-structure Dice for SubCellSeg (DynUNet) against eight baselines including nnU-Net v2, individual DynUNet specialists, Otsu thresholding, Nellie, CellPose v4, StarDist, MicroSAM (ViT-B-LM) and SAM (ViT-B). SubCellSeg achieves the highest Dice on every structure. Both panels share the Dice colour scale (0–1). Off-diagonal mean Dice in (a): 0.50.

### SubCellSeg outperforms other state-of-the-art segmentation methods for 3D segmentation of subcellular structures

To place SubCellSeg in context relative to existing segmentation methods, we compared it against eight baseline methods on the same 13 structures and the same held-out test set: 1) nnU-Net v2 [27] trained as a single foreground/background model on the pooled data of all 13 structures; 2) the structure-specific DynUNet specialist models; 3) Otsu thresholding [9]; 4) Nellie [31], a parameter-free organelle segmentation pipeline designed for 3D live-cell fluorescence microscopy; 5) pretrained StarDist-3D [32, 33] (3D demo model); 6) pretrained CellPose v4 (`cpsam`) [34]; 7) zero-shot Segment Anything (SAM, ViT-B) with automatic mask generation [23], and 8) MicroSAM (ViT-B-LM), a SAM variant fine-tuned on light-microscopy data [24]. Instance segmentation outputs were binarized for a fair Dice comparison. nnU-Net was trained from scratch on the same data seen by SubCellSeg; the other baselines were used zero-shot because they are designed for instance segmentation and cannot be retrained on binary semantic masks without changing what they predict. All methods were evaluated on the same 57 test images per structure (741 in total).

From these experiments, we had three main observations. First, other neural networks (e.g., nnU-Net) could also potentially be used as an alternative backbone of SubCellSeg, similar to our aforementioned comparison of DynUNet, AttentionUNet and SwinUNETR; DynUNet performed slightly better as a backbone than the others. Second, we can observe the advantage of SubCellSeg in 3D subcellular structure segmentation compared with general deep learning models not developed for subcellular structures: CellPose (0.251), StarDist (0.249), MicroSAM (0.032), SAM (0.025) versus SubCellSeg (0.909). We hypothesize that the large variation in the morphologies of different organelles, and in how they are visualized in high-resolution 3D fluorescence microscopy, is the main challenge for these general deep learning models. Third, SubCellSeg, as a deep learning-based generalist model, generalized better than generalist tools and general algorithms based on classic image processing techniques. For example, Nellie, a recent tool among the baselines with the similar goal of segmenting fluorescence images of subcellular structures, did achieve similar performance across different structures, but is on average 0.3 lower in Dice (even more so for Otsu thresholding) than the deep learning-based SubCellSeg.

We also tested structure-specific software on the structures they were developed for (Supplementary Table 4), as well as a state-of-the-art promptable tool. Structure-specific tools, such as ERnet [35] (a dedicated ER segmenter), PlantSeg [36] (developed for cell boundaries) and Spotiflow [37] (a spot detector, applied to the punctate desmosomes and endosomes), did not generalize well to our test set (up to 0.4 lower in Dice than SubCellSeg). The promptable 3D model nnInteractive [38] reached a mean Dice of 0.034 across all 13 structures with a field-of-view box prompt, and 0.036 even with an oracle prompt (the reference mask bounding box plus five corrective clicks).

### Fine-tuning adapts SubCellSeg to new structures with ten times fewer labeled images than training from scratch

To test whether SubCellSeg can be adapted to structures beyond the 13 it was trained on, we evaluated how well its pretrained weights serve as a starting point for new segmentation tasks with application-specific segmentation targets. We performed a controlled fine-tuning study on two structures that were not in the training set: nuclear envelope via lamin B1 and lysosomes via LAMP-1. For nuclear envelope, the morphology is qualitatively similar to structures seen during training (a shell-like boundary around the nucleus). However, when nuclear envelope images are used as a training assay for nuclear segmentation (i.e., extracting the 3D nuclear shapes from nuclear envelopes to serve as the segmentation ground truth for nuclear segmentation from microscopy images of the nuclear dye) [1], the segmentation target becomes a filled shape, instead of a shell. For lysosomes, LAMP-1 localizes to the lysosomal membrane. When our goal is to analyze lysosomes, the segmentation target becomes the filled interior rather than their membrane surface.

We swept training set size (*{*10%, 20%, 50%, 100%*}* of the available labeled images, corresponding to 160 training and 40 validation images for 100% training set) and number of fine-tuning epochs (*{*10, 50, 100, converged*}*), and compared two strategies: (i) full-parameter fine-tuning from the SubCellSeg DynUNet checkpoint, and (ii) training a DynUNet from random initialization with the identical hyperparameter schedule (Fig. 4a and Fig. 4b; for details see the Methods). Out of the box, the base SubCellSeg model reached a Dice score of 0.312 on the nuclear envelope and 0.734 on lysosomes, possibly because lysosomes, like the endosomes seen during training, appear as small puncta. At this resolution, most lysosomes are close to the diffraction limit, so a filled mask and a mask of the punctum are nearly the same, and the filled-volume convention costs little Dice. For the nuclear envelope, the convention matters much more: SubCellSeg segments the envelope as a thin shell, whereas the reference mask is the filled nucleus, which explains the low Dice score. Fine-tuning closed both gaps quickly: with only 10% of labels and 10 epochs, the nuclear envelope model reached a Dice score of 0.699, and with 100% of labels and full convergence, it reached 0.957. Lysosome fine-tuning reached 0.927 at the same setting.

**Figure 4:**
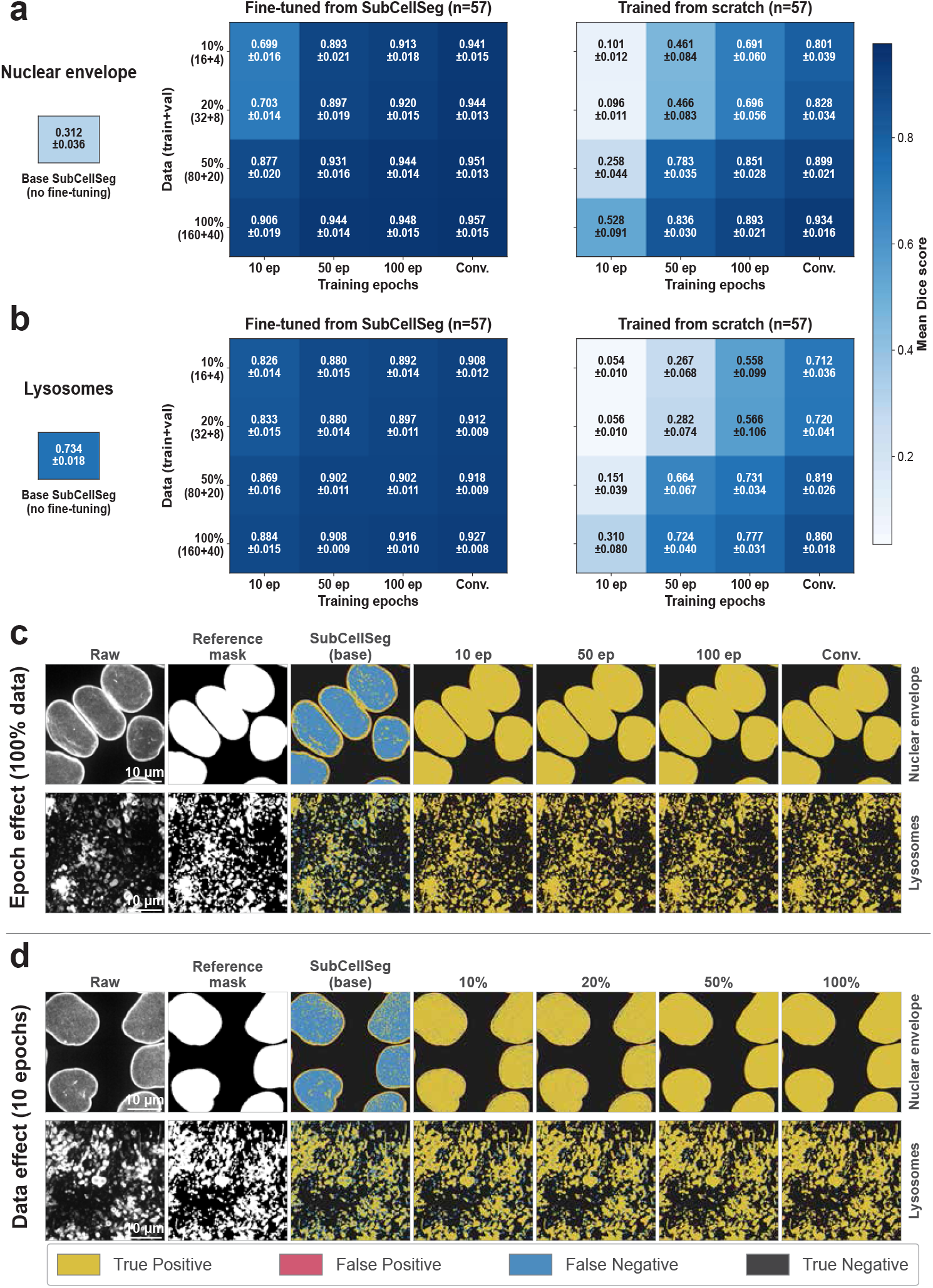
**Data-efficient adaptation to structures outside the training set**. Fine-tuning SubCellSeg on two structures held out of training: nuclear envelope via lamin B1 and lysosomes via LAMP-1, compared against training an identical DynUNet from random initialization under the same hyperparameter schedule. (a) Nuclear envelope. Paired heatmaps show mean Dice on the held-out test set for fine-tuning from the SubCellSeg checkpoint and for training from scratch, as a function of the labeled-data fraction (rows: 10%, 20%, 50% and 100%, corresponding to 16+4, 32+8, 80+20 and 160+40 training + validation volumes) and training length (columns: 10, 50, 100 epochs and converged). Colour encodes mean Dice (scale at right). Applied with no fine-tuning, the base SubCellSeg model reaches Dice 0.312 ± 0.036. Fine-tuning exceeds from-scratch training at every point on the grid, by +0.022 Dice at 100% data and full convergence, the regime most favourable to from-scratch training, and by as much as +0.619 Dice at 50% data and 10 epochs, where the pretrained prior matters most. (b) Lysosomes in the same layout. The base model starts far higher, at Dice 0.734 ± 0.018, and fine-tuning converges to 0.927 ± 0.008. (c, d) Qualitative results for representative test images of the nuclear envelope (top row) and lysosomes (bottom row). Columns show the raw image, the reference mask, the base SubCellSeg prediction before any fine-tuning, and fine-tuned predictions thereafter. (c) Training length at fixed 100% data (10, 50, 100 epochs, converged). (d) Training-set size at fixed 10 epochs (10%, 20%, 50%, 100%). Scale bars, 10 *µ*m.

Fine-tuning SubCellSeg outperformed training from scratch by between +0.022 Dice (100% data, fully converged; the regime where from-scratch training has the most opportunity to catch up) and +0.619 Dice (50% data, 10 epochs; the regime where the pretraining prior matters most; Fig. 4a), across the 4 *×* 4 grid of data fraction *×* training epochs, on the nuclear envelope data. Fine-tuning on 10% of the data (0.941 at convergence) exceeded from-scratch training on 100% of the data (0.934), and the same held for lysosomes (0.908 vs. 0.860; Fig. 4b), a roughly tenfold reduction in the number of labeled images needed. Convergence was also substantially faster: after 100 epochs on 100% of the data, fine-tuning reached 0.948, whereas from-scratch training reached 0.893 (Fig. 4a). In example test images, adding either training epochs (Fig. 4c) or labeled images (Fig. 4d) converged to substantially the same mask.

In practice, 10% of the nuclear envelope training set is 16 labeled volumes, so a few dozen annotated stacks can be enough to deploy SubCellSeg on a previously unseen structure with high accuracy. Lysosomes started from a very different place. Without any fine-tuning, the base model already reached Dice 0.734 *±* 0.018, because most of the places are straightforward for the pretrained model, except the lysosomal membranes need to be segmented as filled to represent lysosomes instead of as shells. Here, fine-tuning lifted this to 0.927 *±* 0.008 at convergence (Fig. 4b). The two structures therefore represent two different scenarios that future users can use as examples while fine-tuning on new tasks when the pretrained SubCellSeg cannot provide satisfactory results.

### SubCellSeg performance is robust to remodeling of subcellular structures in response to drug perturbations

A common concern with segmentation workflows designed for a specific subcellular structure morphology is that they fail when the structure deviates from normal [1]. To evaluate SubCellSeg’s ability to accurately segment subcellular structures as they remodel in response to drug perturbation, we assembled a drug perturbation dataset of 3D images of WTC-11 hiPS cell lines expressing endogenously tagged markers of five structures: mitochondria via TOM20, Golgi via sialyltransferase 1, nuclear pores via NUP153, nucleoli (DFC) via fibrillarin, and microtubules via *α*-tubulin. The nucleoli DFC images (control and actinomycin D) are from a published dataset [3]; the other four lines were imaged for this study following the same protocol (Methods). Drug-treated cells were imaged alongside vehicle (DMSO)-treated controls. The fluorescently labeled mitochondria and Golgi cell lines were treated with rotenone, an inhibitor of mitochondrial complex I [39] that fragments the mitochondrial network, depletes cellular ATP, which in turn disassembles the cis-Golgi network [40], and compromises microtubule integrity through direct tubulin binding [41, 42]. The nuclear pore line was treated with thapsigargin, a SERCA inhibitor that depletes ER calcium stores [43]. The nucleoli DFC line was perturbed with actinomycin D, which at low doses selectively inhibits RNA polymerase I and reorganizes nucleolar components into caps [44], and the microtubule line was treated with nocodazole, which at nanomolar concentrations blocks cells in mitosis without net microtubule disassembly by reducing microtubule turnover [45]. For each condition, we benchmarked SubCellSeg against the corresponding structure-specific specialist model on the same images.

Regardless of the treatment conditions or the structure analyzed, SubCellSeg produced biologically plausible segmentations consistent with the phenotypic changes expected for each perturbation (Fig. 5). In the mitochondria line, both SubCellSeg and the specialist model recovered the fragmented mitochondrial pool following rotenone treatment (Fig. 5e). This was the one condition in which all methods remained robust. The Golgi line proved more discriminating (Fig. 5d): SubCellSeg accurately captured the Golgi-dispersal phenotype at high rotenone concentrations, whereas the others suffer from over-segmentation. Furthermore, SubCellSeg accurately recognized the redistribution of nucleoporins into concentrated foci in the nuclear pore line under thapsigargin (Fig. 5b), the disassembly of the nucleoli DFC under actinomycin D (Fig. 5a), and the mitotic microtubule reorganization under nocodazole (Fig. 5c). Taken together, these results establish a clear evidence of robustness, with SubCellSeg outperforming the specialist models and classic image segmentation workflows.

**Figure 5:**
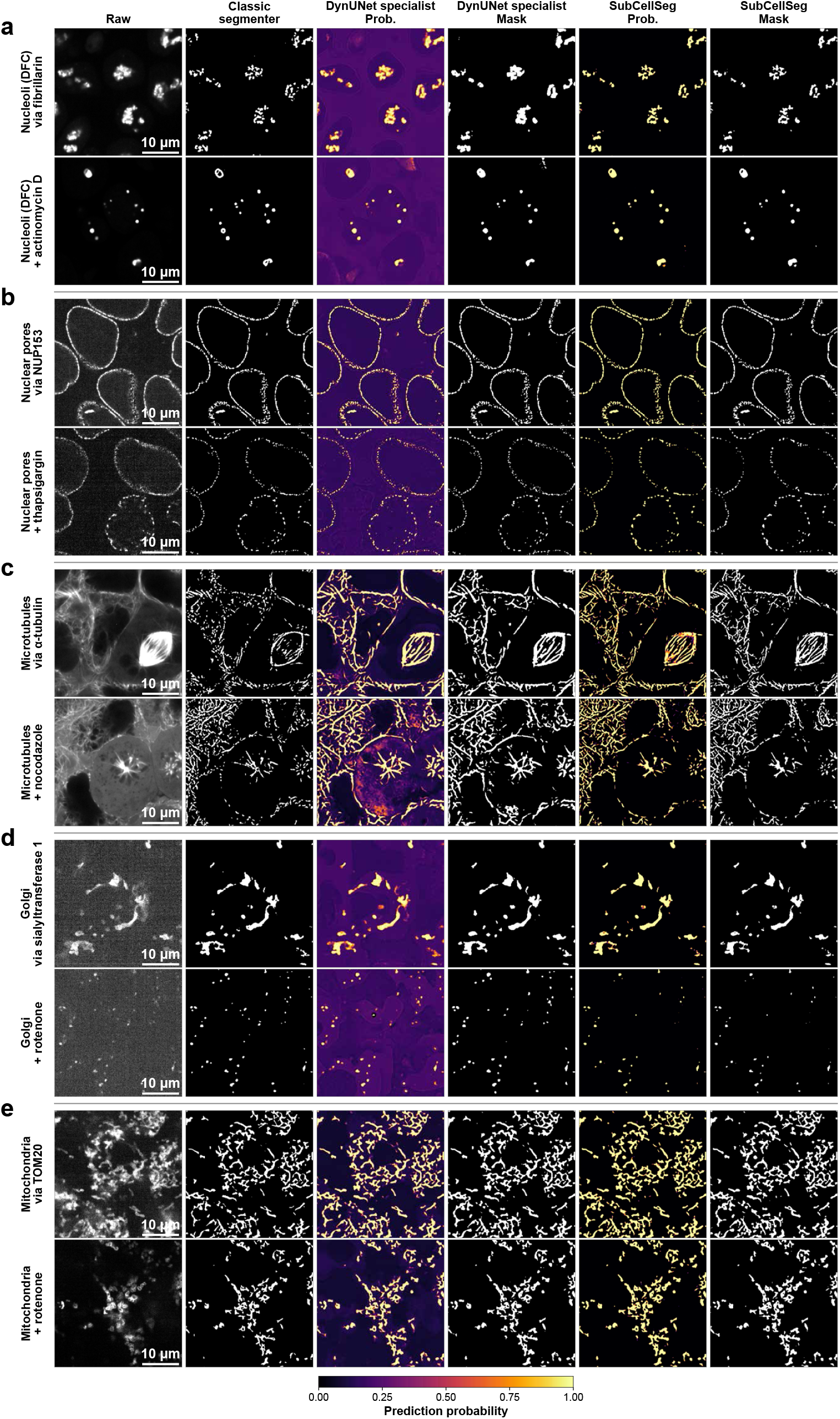
**Robust segmentation under drug-induced structure remodeling**. Live WTC-11 hiPS cells expressing endogenously tagged markers of five structures, imaged in 3D on the training microscope (SDCM-1, 100*×*) after pharmacological perturbation, alongside vehicle (DMSO) controls; acquisition settings in Methods. For each condition, columns show the raw image, the results from classic image segmentation workflows, the structure-specific specialist probability map and binary mask, and the SubCellSeg probability map and binary mask. Probability maps use the Inferno colour scale (prediction probability 0–1). (a) Nucleoli DFC via fibrillarin under actinomycin D, which at low doses selectively inhibits RNA polymerase I and reorganizes nucleolar components into caps; (b) Nuclear pores via NUP153 under thapsigargin, which depletes ER calcium stores and induces ER stress; nucleoporins redistribute into concentrated foci; (c) Microtubules via *α*-tubulin under nocodazole, which at nanomolar concentrations blocks mitosis without net microtubule disassembly; (d) Golgi via sialyltransferase 1 under rotenone. (e) Mitochondria via TOM20 under rotenone, an inhibitor of mitochondrial complex I that fragments the mitochondrial network; the one condition in which all methods, including the classic Segmenter, remained robust. Nuclear pores, Golgi and mitochondria were part of SubCellSeg’s training set whereas nucleoli DFC and microtubules were not, and neither SubCellSeg nor the specialists saw any perturbed cell during training. SubCellSeg tracks the expected phenotype in every case, while the specialists are less consistent. Scale bars, 10 *µ*m.

### SubCellSeg generalizes out of the box to other microscopes, magnifications and cell types

Another important aspect to evaluate was whether SubCellSeg overfits to the specific point-spread function, signal-to-noise, and voxel resolution of the spinning-disk confocal system or cell type used for training. If it does, SubCellSeg would be much less accurate on images acquired with other microscopes or objectives, or of other cell types. We, therefore, evaluated the base SubCellSeg model visually (no reference masks exist for these images), without fine-tuning, on four out-of-distribution test conditions that vary the microscope, magnification, structure, or cell type relative to the training distribution (Fig. 6a).

**Figure 6:**
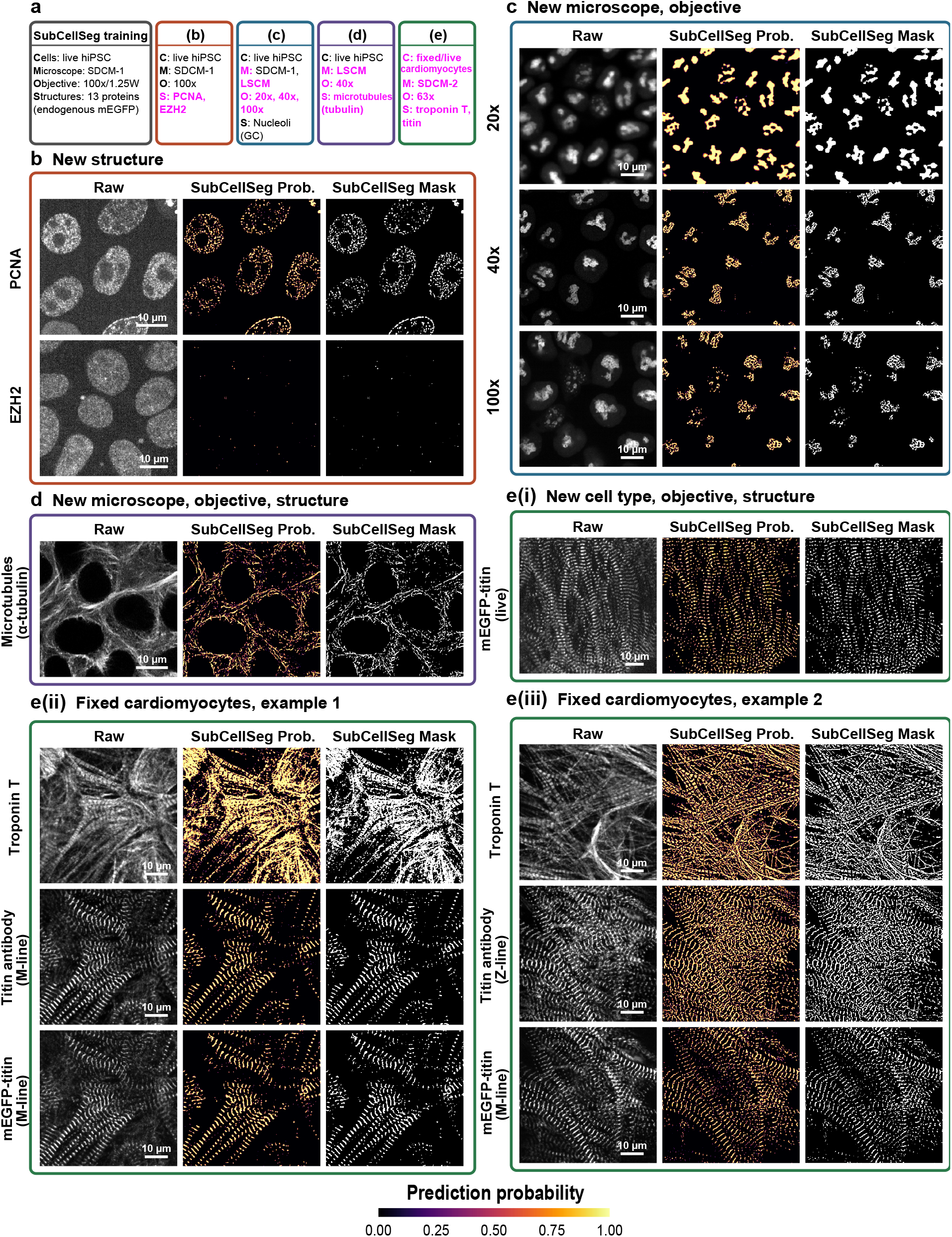
**Zero-shot generalization across microscopes, magnifications, cell types and structures**. SubCellSeg applied with no retraining or fine-tuning. Throughout, columns show the raw fluorescence image, the SubCellSeg probability map and the binarized mask; probability maps share the colour scale at bottom (0–1). (**a**) The four test conditions, each summarized by cells (C), microscope (M), objective (O) and structures (S). Top, the training distribution: live WTC-11 hiPS cells on a spinning-disk confocal microscope (SDCM-1) at 100*×*/1.25 W, 13 endogenously tagged mEGFP structures. Below, conditions **b**–**e**; entries matching training in black, shifted entries in magenta. (**b**) New structures, training instrument: PCNA (replication foci) and EZH2 (diffuse chromatin-associated signal) in live hiPSC on SDCM-1 at 100*×*. (**c**) New microscope and objectives: nucleoli (GC) via nucleophosmin at 20*×* and 100*×* on SDCM-1 and at 40*×* on an Airyscan laser-scanning confocal microscope (LSCM). (**d**) New microscope, objective and structure: microtubules via *α*-tubulin on the LSCM at 40*×*. (**e**) New cell type, objectives and structures in hiPSC-derived cardiomyocytes, with a new microscope in (**ii**, **iii**): sarcomeres. (**i**) Live cells expressing mEGFP-tagged titin imaged on SDCM-1 at 40*×*. (**ii**, **iii**) Fixed cells imaged on a second spinning-disk confocal microscope (SDCM-2) at 63*×*, three rows each: cardiac troponin T (thin filaments); a titin antibody marking the M-line in (**ii**) and the Z-line in (**iii**); and mEGFP-tagged titin at the M-line. Scale bars, 10 *µ*m.

**Figure 7:**
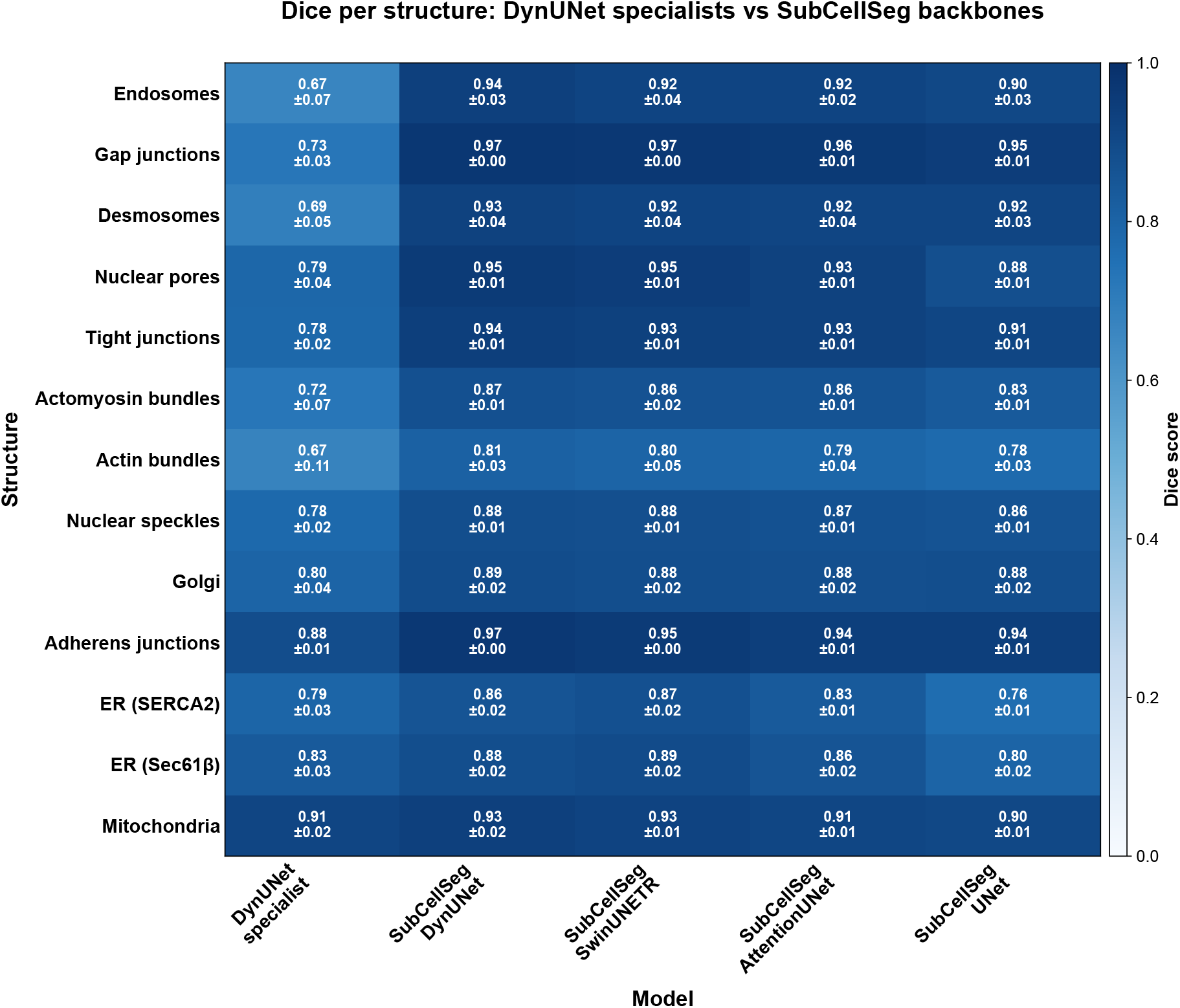
Per-structure Dice for specialists and the four SubCellSeg backbones. Mean Dice over the 57 held-out test images per structure for the structure-specific DynUNet specialists and SubCellSeg trained with DynUNet, SwinUNETR, AttentionUNet and UNet backbones. DynUNet and SwinUNETR outperform the specialists on every structure; the standard UNet falls below the specialists on ER via SERCA2, ER via Sec61*β*, and mitochondria via TOM20 (by 0.01 *−* 0.03), and AttentionUNet ties them on mitochondria. Across the 13 structures, the means are 0.909 (DynUNet), 0.904 (SwinUNETR), 0.892 (AttentionUNet), 0.870 (UNet) and 0.773 (specialists). Backbone choice matters most for ER via SERCA2, ER via Sec61*β*, and nuclear pores via NUP153 (spread 0.08–0.11 Dice, driven by the standard UNet; SwinUNETR is best on both ER markers) and least for Golgi via sialyltransferase 1 and desmosomes via desmoplakin (spread *≤* 0.011).

#### Same microscope, new structure (PCNA & EZH2; SDCM-1 100×)

PCNA and EZH2 were imaged on the same microscope as the training data, so only the structure itself differs from training. PCNA marks DNA replication foci [46], which assemble and disassemble as replication proceeds, so a single marker looks quite different at different cell-cycle stages, and none of these patterns resembles a training structure. EZH2, an enzymatic subunit of Polycomb repressive complex 2 that methylates histone H3 at Lys27 [47], gives a diffuse signal that fills the nucleus without discrete objects or sharp boundaries. SubCellSeg segmented PCNA foci across cell-cycle phases (Fig. 6b, top), but returned only sparse, isolated detections for EZH2 and did not capture its texture (Fig. 6b, bottom). SubCellSeg therefore handles new arrangements of distinct objects, but not signals defined by texture alone.

#### Multiple magnifications and confocal microscopes (nucleoli GC via nucleophosmin; SDCM-1 20×, LSCM 40×, SDCM-1 100×)

The nucleolar granular component, marked by nucleophosmin, is a low-contrast nuclear compartment that was not part of training. SubCellSeg returned nucleolar masks at all three magnifications (Fig. 6c), with quality scaling with effective resolution.

*New microscope, new structure (microtubules via mEGFP-tagged α-tubulin; Airyscan LSCM* 40*×)*

Tubulin was not in the training set, and the Zeiss Airyscan provides higher lateral resolution but a different PSF and pixel size. SubCellSeg produced consistent tubulin segmentations (Fig. 6d), capturing microtubule density and spatial distribution.

#### New cell type, new magnification, and new microscope (cardiomyocyte sarcomeres; SDCM-1 40× and SDCM-2 63×)

hiPS cell-derived cardiomyocytes expressing mEGFP-tagged titin [48] differ from the training set in both structure and cellular context. SubCellSeg segmented the striated sarcomere pattern and preserved the alternating banding in both fixed and live imaging (Fig. 6e), demonstrating that SubCellSeg generalizes beyond hiPS cells.

A further set of zero-shot evaluations was performed across 12 additional structures from the WTC-11 dataset that were not part of training (plasma membrane via CAAX, actin filaments via *β*-actin, centrioles via centrin-2, nucleoli (DFC) via fibrillarin, histones via H2B, lysosomes via LAMP-1, nuclear envelope via lamin B1, nucleoli (GC) via nucleophosmin, matrix adhesions via paxillin, peroxisomes via PMP34, cohesins via SMC-1A, microtubules via *α*-tubulin). Performance was variable (Extended Data Fig. 2) but consistently good for structures with high-contrast morphologies (centrioles, nucleoli (DFC), nucleoli (GC), matrix adhesions and microtubules), and weaker for structures whose reference masks follow conventions that differ substantially from training (plasma membrane, actin filaments, lysosomes and nuclear envelope).

## Discussion

A single model, SubCellSeg, trained jointly on paired fluorescence and reference mask images of 13 subcellular structures, segmented each of them more accurately than specialist models trained on each structure alone (Fig. 1). This suggests that training across a strategically selected collection of structures with consistent imaging parameters and curated reference masks [1] lets the model learn what fluorescence images of subcellular structures have in common, such as how signal separates from background and out-of-focus light, rather than features tied to the shape of one structure. The cross-evaluation (Fig. 3a) supports this interpretation. Each specialist model exhibits high accuracy on the structure it was trained on but degrades sharply when applied to other structures, indicating that its learned features are narrowly structure-specific. Further, for subcellular structures with similar morphology and phenotype (e.g., the specialist for ER via SERCA2 evaluated on ER via Sec61*β*), specialist models perform better compared to structures with significant differences in appearance, fluorescence signal profile, and distribution (the specialist for ER via SERCA2 evaluated on desmosomes and endosomes). In contrast, SubCellSeg maintains high accuracy across all structures with a single set of weights, reflecting learned, reusable image features for fluorescence contrast that generalize across structures (shown in Fig. 3b). Consistent with this, clustering the cross-evaluation matrix splits the structures into two groups by image statistics rather than biological category, separating spatially extended, high-foreground-fraction structures from sparse, low-foreground-fraction ones (an *∼*8.5-fold difference in mean foreground occupancy); notably the split cuts across biology, with adherens junctions grouping among the dense structures and tight junctions among the sparse ones, indicating that the specialist features organize by fluorescence contrast rather than structure identity.

The architecture comparison (Fig. 1c) further reinforces the benefit of multi-structure training. All four SubCellSeg backbones exceeded the DynUNet specialists, and the spread between backbones (*∼*4 Dice points from DynUNet to UNet) was smaller than the gap between the specialists and SubCellSeg DynUNet (*∼*14 Dice points). This indicates that, within the architectures tested, training strategy had a larger effect on accuracy than the backbone choice. Training-set diversity also matters: in an ablation over 2–14 training structures, in-distribution Dice rose with the number of structures, but generalization to held-out structures was non-monotonic, and adding plasma membrane via CAAX degraded the other structures (Extended Data Fig. 3).

The fine-tuning results indicate that SubCellSeg can serve as a pretrained representation for subcellular structure segmentation (a foundation model [25]) that transfers to new and previously unseen structures in 3D fluorescence microscopy imaging. For both the nuclear envelope and lysosomes, fine-tuning on 10% of the labels outperformed training from scratch on all of them (Fig. 4), which points to a practical workflow. Rather than training a deep model from scratch for a new structure, one could collect a few dozen labeled volumes and fine-tune from SubCellSeg. The advantage is largest at small training set sizes and narrows as more data is available, consistent with prior transfer-learning studies [49, 50].

Structure morphology also changes within a single experiment: pharmacological or genetic perturbations remodel structure shape on minute-to-hour timescales. We asked whether SubCellSeg, which was trained only on untreated WTC-11 cells, can segment perturbed structures at all, and whether it tracks the morphological changes faithfully. The drug-perturbation experiment addresses this directly. SubCellSeg saw healthy mitochondria and Golgi during training but never fragmented mitochondrial networks or dispersed Golgi. That its masks remained visually consistent with the perturbed morphologies suggests it remains useful when structure morphology shifts away from the untreated training examples. The nucleoli DFC and microtubule cell lines make an even stronger case. SubCellSeg was never exposed to these structures during training in either the untreated or the perturbed condition, yet its masks followed both structures closely, suggesting that its learned features generalize to entirely unseen structures as well as to unseen morphologies. This supports using SubCellSeg in perturbation experiments. This matters because almost every biologically meaningful experiment involves perturbing the system, and a model that fails when morphology changes is of limited use for biological research. Such failures also force biologists to switch between different models when segmenting the same structure across drug conditions, leading to inconsistent, non-standardized analyses. SubCellSeg instead provides a single framework that segments structures across both perturbed and control conditions.

SubCellSeg performance was also found to be robust to change in acquisition settings. Without any fine-tuning, SubCellSeg produced reasonable segmentations on data from different microscopes, magnifications, and cell types, i.e., microtubules on an Airyscan system, nucleoli-GC across a 20*×*–100*×* range, and sarcomeres in iPSC-derived cardiomyocytes imaged both fixed and live (Fig. 6). That the model transfers to a cell type and structures well outside the training distribution indicates that it relies on image content rather than instrument-specific features. The generalizability of SubCellSeg is not unbounded though. EZH2, a diffuse chromatin-associated signal with no sharp object boundaries, marks a current limit: structures defined by texture rather than by distinct morphology remain hard, and no model can recover detail that the optics did not resolve in the first place. Perhaps with structure specific fine-tuning, versions of SubCellSeg could be trained that can overcome this challenge.

Several limitations deserve explicit mention. First, all training data comes from a single cell line (WTC-11 hiPSC), generated in a single laboratory and predominantly using a single instrument configuration. The cross-microscope and cardiomyocyte results suggest that the model generalizes beyond the standard setting. But this evaluation is limited to the data considered here, and its breadth is not sufficient to claim general coverage. Second, SubCellSeg performs semantic segmentation. For analyses that require individual structure counts (e.g., per-cell mitochondrion number), SubCellSeg masks must be processed downstream, e.g., with watershed-based instance extraction. This setup was intentional, as an instance is not well defined for all structures and is heavily context-dependent. Finally, today’s foundation models for biological imaging do not address the 3D subcellular fluorescence regime. MicroSAM [24], despite being fine-tuned on light-microscopy data, reached only a 0.032 Dice score (zero-shot; Table 2); its EM-structure variants scored near zero, reflecting the modality gap between electron and fluorescence microscopy. MedSAM [51], trained on clinical imaging with bounding-box prompts, targets a different interaction paradigm and modality and was therefore excluded from the zero-shot comparison. Building a foundation model directly on subcellular fluorescence structures is an appealing next step that SubCellSeg’s dataset could support.

**Table 2:** Comparison to state-of-the-art segmentation methods. Mean Dice score across 13 trained structures on the held-out test set (57 test images per structure, 741 total). All methods evaluated on identical samples. “Trained” indicates the method was trained on the same data SubCellSeg saw; “zero-shot” indicates the method was used with publicly available pretrained weights, with instance outputs binarized for Dice computation. Statistical comparison: paired Wilcoxon signed-rank test, SubCellSeg DynUNet vs. each baseline, *p <* 10*^−^*^7^ for every per-structure comparison. Per-structure values in Supplementary Table 2.

| Method | Mode | Mean Dice (n=13) |
|---|---|---|
| **SubCellSeg (DynUNet)** | Trained, multi-structure | **0.909** |
| nnU-Net v2 [27] | Trained, multi-structure | 0.841 |
| Individual DynUNet specialists | Trained, per-structure (×13) | 0.773 |
| Nellie [31] | Zero-shot (no learning, organelle-specific) | 0.600 |
| Otsu thresholding [9] | Zero-shot (no learning) | 0.350 |
| CellPose v4 [34] | Zero-shot (pretrained) | 0.251 |
| StarDist-3D (3D demo) [33] | Zero-shot (pretrained) | 0.249 |
| MicroSAM (ViT-B-LM) [24] | Zero-shot (LM fine-tuned, 2D) | 0.032 |
| SAM (ViT-B, AMG) [23] | Zero-shot (pretrained, 2D) | 0.025 |

## Methods

### Data sources

#### Cell lines

Human induced pluripotent stem (hiPS) cell lines were generated by the Allen Institute through endogenous tagging of marker proteins [26, 48]. All but AICS-0007 cl.79 ACTN1-mEGFP (the alpha-actinin-1 line) [26] are available through the Allen Cell Collection (https://www.allencell.org), along with the protocols for hiPS cell culture and cardiomyocyte differentiation. Supplementary Table 5 lists the clone, tagged protein, structure, and use of each line in this study.

#### Reference imaging configuration

Images of the 13 cell lines used for training and of the 12 unseen-structure lines (Supplementary Table 5) were taken from the WTC-11 hiPSC Single-Cell Image Dataset v1 [2]. The settings and configuration of the spinning-disk confocal microscope (SDCM-1) used to collect these data are described in [2] and will be referred to here as the Reference imaging configuration. Only the channel corresponding to the tagged protein was used in this study.

#### Drug perturbation imaging

The drug perturbation dataset (Fig. 5) combines two images from [3], an untreated control and cells treated with actinomycin D for 4 h, both of AICS-0014 (the fibrillarin line), with images acquired for this study. Following the cell handling and drug administration procedures described in that study, additional images were acquired using AICS-0011 (the TOM20 line) and AICS-0025 (the sialyltransferase 1 line) treated with 0.4 *µ*M rotenone (MedChemExpress, HY-B1756) for 24 h, AICS-0069 (the NUP153 line) treated with 1 *µ*M thapsigargin (MilliporeSigma, SML1845) for 4 h, and AICS-0012 (the *α*-tubulin line) treated with 0.1 *µ*M nocodazole (MedChemExpress, HY-13520) for 2 h. DMSO-treated controls were also imaged. All images were acquired as described in [3], with three differences from the Reference imaging configuration: exposure time was reduced to 100 ms, 488-nm laser power was doubled (to approximately 4.6 mW), and only the 488-nm fluorescence and transmitted-light channels were acquired, as no additional dyes were used.

#### Zero-shot inference imaging

The images in Fig. 6b–d were not used for training, hyperparameter tuning, or checkpoint selection. High-magnification (100*×*) images of AICS-0088 (the PCNA line), AICS-0095 (the EZH2 line), and AICS-0057 (the nucleophosmin line) were produced using the same modifications to the Reference imaging configuration described for the drug perturbation imaging, except for laser power (see Supplementary Table 6). AICS-0057 (the nucleophosmin line) was also imaged using the same configuration but with a 20*×*/0.8 NA Plan-Apochromat objective and no binning. AICS-0057 (nucleophosmin) and AICS-0012 (α-tubulin) lines were imaged on a laser scanning confocal microscope (LSCM; ZEISS LSM 880 with an Airyscan detector and a C-Apochromat 40*×*/1.2 W Korr FCS M27 objective). These images were acquired using Fast Airyscan in Opt mode with an acquisition frame size of 804 *×* 804 pixels. Channels were acquired line-sequentially using 405-, 488-, and 633-nm excitation. Fluorescence was detected with the Airyscan detector at a detector gain of 900 for *α*-tubulin and 850 for nucleophosmin, and a digital gain of 1.0. Images were processed as 3D image stacks using 3D Airyscan processing in ZEISS ZEN 2.3 (Black edition) with automatic processing strength, changing the x-y image dimensions from 804 *×* 804 to 792 *×* 792 pixels. Cells were maintained at 37 *^◦^*C with 5% CO_2_ in a humidified environmental chamber during imaging. Cells were labeled with NucBlue Live (Thermo Fisher Scientific) and CellMask Deep Red (CMDR; Thermo Fisher Scientific) as described in [2]. Additionally, in the case of imaging of AICS-0012 ( the *α*-tubulin line), a photoprotective cocktail consisting of 1 mM ascorbic acid, 0.3 U/ml OxyFluor (Oxyrase Inc.) and 10 mM lactate was used. Only the 488-nm channel was used for the present study. Supplementary Table 6 lists pixel dwell time, laser power, and voxel size for each panel. Images were passed to SubCellSeg at their native voxel size, without resampling.

#### Cardiomyocyte imaging

Live cardiomyocytes (Fig. 6ei) were imaged on the SDCM-1 system with modification to the Reference imaging configuration where the objective, exposure times, and laser powers were changed (see Supplementary Table 6). Fixed cells (Fig. 6eii and eiii)were imaged on a different spinning-disk confocal microscope (SDCM-2); imaging and immunolabeling were performed as described in[52]. Primary antibodies used were mouse anti-cardiac troponin T (Thermo Fisher Scientific, MA5-12960) and rabbit anti-titin, either against the C-terminus to mark the M-line (M8–M10; MyoMedix, TTN-9) or against Z1/Z2 to mark the Z-line (MyoMedix, TTN-1). Secondary antibodies were goat anti-mouse Alexa Fluor 647 (A-21235) and goat anti-rabbit Alexa Fluor 568 (A-11011; both Thermo Fisher Scientific). All antibodies were used at 1:1000, and nuclei were counterstained with DAPI. The troponin T, titin-antibody and mEGFP-titin channels were each segmented; the nuclear and membrane channels were not used. These images were likewise not used for training or model selection.

#### Reference segmentations

Reference segmentation masks for all 13 training structures were generated using the Allen Cell and Structure Segmenter [1], a classical image-processing toolkit that combines structure-specific intensity thresholding, morphological filtering, and topology-aware operations. These are not ground truth: each structure-specific workflow was evaluated per structure, and only structures whose reference segmentations were judged sufficiently accurate were retained (Results). The retained reference segmentations were then treated as pseudo-ground truth and used as the segmentation target for training and validation, and as the reference masks for evaluation. Reference segmentation masks for the fine-tuning structures (nuclear envelope via lamin B1 and lysosomes via LAMP-1) were generated in the same way with their structure-specific workflows. For images outside the WTC-11 hiPSC Single-Cell Image Dataset v1, reference segmentation masks were generated with the structure-specific workflow for the target structure.

#### Data splits

For each structure, images were split into training (160 z-stacks, *∼*60%), validation (40 z-stacks, *∼*15%), and test (67 z-stacks, *∼*25%) sets with no field-of-view overlap. Splits were stratified to ensure representation of diverse imaging sessions and passage numbers. All benchmark numbers reported here are computed on the held-out test sets. During evaluation, a small number of test images showed unusually low segmentation accuracy. Examining these images revealed that their reference masks, generated by the Allen Cell and Structure Segmenter, captured only part of the visible structure. The problem was most pronounced for desmosomes via desmoplakin, where biologists confirmed the deficiency in the reference workflow, and similar but less frequent cases were found for other structures (Extended Data Fig. 4). To treat all structures equally, we set the number of images to exclude by the most affected structure (10 images, for desmoplakin) and removed the 10 test images with the lowest SubCellSeg DynUNet Dice from every structure. The excluded set was largely the same regardless of which backbone was used for ranking (mean pairwise overlap *∼*90% across the four backbones), and the same images were excluded for all methods and analyses, leaving 57 z-stacks per structure (741 in total). Our conclusions do not depend on this step: on the complete test set of 67 z-stacks per structure, SubCellSeg exceeds the specialist models by 0.124 Dice, compared with 0.136 after curation (Supplementary Note 1, Supplementary Table 1). The excluded images, their reference masks, all model predictions, and per-image scores are provided in the SubCellSeg dataset.

#### Model architectures

All SubCellSeg models were implemented using the MONAI [53] and cyto-dl frameworks [54]. The four backbones evaluated were:

- **DynUNet** [27, 53]: MONAI’s dynamic U-Net, adapted from the nnU-Net architecture. Unlike nnU-Net, kernel sizes, strides, and patch size are specified directly rather than inferred by a self-configuring planner; we used 32 *×* 128 *×* 128 patches with strides [1, 2, 2, 2, 2] and 3 *×* 3 *×* 3 kernels throughout.
- **AttentionUNet** [29]: A 3D U-Net with additive attention gates on each skip connection, which reweight encoder features before they are concatenated into the decoder. Channels (16, 32, 64, 128, 256) and strides (2, 2, 2, 2).
- **SwinUNETR** [28]: A hierarchical Swin Transformer encoder with a CNN decoder, using the feature size 48 (twice MONAI’s default of 24) and depths (2, 2, 2, 2).
- **UNet** [30, 55]: MONAI’s residual 3D U-Net with channels (16, 32, 64, 128, 256), strides (2, 2, 2, 2) and two residual units per resolution level.

All models take a single-channel 3D patch as input and output a single-channel probability map of the same spatial dimensions. No structure-specific input information (e.g., one-hot encoding, structure ID, FiLM conditioning) is provided at training or inference time.

### Training protocol

#### Multi-structure training

SubCellSeg was trained jointly on all 13 structures by interleaving batches from different structures within each epoch. Each batch draws patches from images sampled at random from the union of all 13 training sets; because every structure contributes the same number of images (160), structures are sampled uniformly in expectation and batches routinely mix structures. The loss used for all model training is the Generalized Dice Focal Loss, a combination of generalized Dice loss [56, 57] and focal cross-entropy loss [58] as implemented in MONAI [53], which helps balance foreground–background class imbalance. Class imbalance varies substantially across structures (desmosomes have very sparse foreground; ER via Sec61*β* fills much of the cell volume), and the focal term is needed to prevent the loss from being dominated by easy structures.

#### Hyperparameters

Models were trained for 1000 epochs on a single NVIDIA A100 GPU using PyTorch with mixed precision. We used Adam [59] with a learning rate of 10*^−^*^4^ and weight decay of 10*^−^*^4^, decaying the learning rate exponentially by a factor of 0.995 per epoch. Each step used 32 patches of 32 *×* 128 *×* 128 voxels, drawn one per image by random multi-scale cropping. Raw intensities were z-score normalized channel-wise. Augmentation was applied to the raw channel only and consisted of random histogram shift, random standard-deviation intensity shift (factor 0.1), and random contrast adjustment (*γ ∈* [0.9, 1.5]), each applied with probability 0.1. Validation ran every epoch; the checkpoint with the lowest validation loss was retained, and training stopped early if validation loss did not improve for 100 consecutive epochs.

#### Specialist baselines

Structure-specific specialist DynUNet models were trained using the identical con-figuration (patch size, batch size, learning rate, optimizer, augmentation, epochs) as SubCellSeg, with the only difference being that each was trained on images from a single structure. Both were trained under the same early-stopping rule (no improvement in validation loss for 100 epochs), so each specialist was trained to convergence on its own data, and performance differences reflect the multi-structure training data rather than an unequal number of training epochs. Specialists were trained with the DynUNet backbone only, the default SubCellSeg architecture, so that each specialist and SubCellSeg differ only in their training data.

#### Fine-tuning protocol

For Fig. 4, we initialized model weights from the pretrained SubCellSeg DynUNet checkpoint and updated all parameters (full fine-tuning). The fine-tuning learning rate was 5 *×* 10*^−^*^5^ (half of the from-scratch rate); we observed empirically that lower learning rates yielded slightly better data efficiency and final performance. From-scratch baselines used the standard 10*^−^*^4^ learning rate. Both strategies used the same augmentation pipeline and loss function. Training subsets of *{*10%, 20%, 50%, 100%*}* of available labels were constructed by stratified random sampling from the available nuclear envelope and lysosomes annotations. In every case, the dataset scaling is only applied to the train and validation sets (e.g., 20% of train data (total 160) = 32 and 20% of validation data (total 40) = 8), while the test set remains unchanged with 57 z-stacks. Each fine-tuning configuration was run for *{*10, 50, 100*}* epochs, plus a ”converged” condition trained until validation loss did not improve for 50 consecutive epochs.

#### Cross-evaluation experiment

For Fig. 3a, each of the 13 specialist DynUNet models was applied to all 13 test sets, yielding a 13 *×* 13 Dice matrix. The diagonal of this matrix corresponds to a specialist applied to its target structure. The off-diagonal entries correspond to a specialist applied to a structure it was not trained on.

#### State-of-the-art comparison

For Fig. 3b and Table 2, eight baseline methods were evaluated on the same 13 trained structures and the same held-out test set as SubCellSeg. **nnU-Net v2** [27] was trained with the `3d fullres` configuration as a single binary (foreground/background) task on the pooled data of all 13 structures (single fold, default pre-processing). **Per-structure DynUNet specialists** are the same specialists used in the main benchmark. **Otsu thresholding** was applied per volume to the raw intensity with no pre-processing. **Nellie** [31] was run zero-shot with all default parameters. We only used its segmentation stages (Filter, then Label) since the tracking and feature-extraction stages do not apply to single-timepoint masks. Because Nellie’s filter scales are defined in physical units, the axis order (ZYX) and voxel size of the source acquisitions (0.29 *×* 0.108 *×* 0.108 *µ*m) were explicitly passed to Nellie. **StarDist-3D** [33] was used zero-shot with the `3D demo` pretrained model. **CellPose v4** (`cpsam`) [34] was used zero-shot with 3D mode (auto-diameter, single channel). **Segment Anything** [23] (ViT-B) and **MicroSAM** [24] (ViT-B-LM) were applied with automatic mask generation per 2D slice with no z-consistency. Instance outputs from CellPose, StarDist, and SAM and Nellie were binarized (`label map > 0`) before Dice computation. The same test set of 57 images per structure is used (n=741 total). Statistical comparison used paired Wilcoxon signed-rank tests on per-sample Dice scores.

#### Structure-specific and promptable baselines

Four further tools were evaluated zero-shot, with their released weights and no fine-tuning, on the structures matching their design and on the same 57 test images per structure. Each input was calibrated to the tool’s expected voxel size, using our voxel size of 0.29 *×* 0.108 *×* 0.108 *µ*m (*z × y × x*). ERnet [35] (single-frame SwinIR model, `20220306 ER 4class swinir nch1`) was applied slice-wise to ER via SERCA2 and ER via Sec61*β*, following ERnet’s own stack pipeline (per-slice 1–99th-percentile normalisation). Slices were upsampled to its 42 nm training pixel size, and the binary prediction was downsampled back. Foreground was any non-background class (tubule, sheet or sheet-based tubule). PlantSeg [36] (`generic confocal 3D unet`) was applied to adherens junctions via *β*-catenin, the one membrane-like structure, after rescaling the volume to the model’s 0.235 *×* 0.150 *×* 0.150 *µ*m voxel size. The boundary probability was resampled back and thresholded at 0.5. Spotiflow [37] (`smfish 3d` and `synth 3d`, default probability threshold) was applied to the punctate desmosomes via desmoplakin and endosomes via RAB5A after resampling to isotropic 0.108 *µ*m voxels. Because Spotiflow outputs points rather than masks, Spotiflow and SubCellSeg were compared on DSP and RAB5A by object-level detection F1 rather than voxel-wise Dice. Reference points and SubCellSeg points were the centroids of connected components of the reference mask and SubCellSeg binary masks, respectively, and points were matched one-to-one within 1 *µ*m (Hungarian matching; results at 0.5 *µ*m differed by at most 0.012). nnInteractive [38] (v1.0) was applied to all 13 structures on the raw volume, without normalisation or resampling, as its documentation recommends. Each structure was treated as a single target, using two prompting protocols. The label-free protocol placed one 2D box spanning the full field of view on the brightest *z*-slice. The oracle protocol used the reference mask bounding box on the *z*-slice with the largest reference mask area, followed by five corrective clicks, each placed at the deepest voxel of the largest error component. Paired comparisons with SubCellSeg used two-sided Wilcoxon signed-rank tests on per-image scores.

Several other structure-specific tools were considered but not run, because their input domain or output type does not match 3D binary masks of the trained structures. For the ER, nERdy and nERdy+ [60] and MRE [61] segment 2D ER (and, for MRE, mitochondrial) network images. For mitochondria, MitoGraph [13] reconstructs 3D surfaces and skeletons, Mitochondria Analyzer [62] and MiNA [63] are Fiji threshold-and-skeleton workflows built primarily for morphology quantification, and MitoSegNet [64] is a 2D model trained on *C. elegans* muscle. For filaments, SOAX [65] extracts 3D centerlines rather than volumes. CellSAM [66] targets 2D whole-cell segmentation. The largest multi-organelle benchmark, OpenOrganelle [67], is built on volume electron microscopy. Related multi-organelle work in live-cell fluorescence segments organelles from a single lipid-dye channel [68], a different input modality from tagged-protein stacks. We found no released specialist segmenters for the Golgi, nuclear speckles, tight junctions or gap junctions.

#### Probability map thresholding

SubCellSeg outputs a probability map: a floating-point volume where each voxel value represents the model’s confidence that the voxel belongs to the foreground. To convert to a binary mask, we apply a fixed threshold of 0.5 (corresponding to threshold 128 when the probability map is stored as a uint8 image). A fixed threshold is sufficient because SubCellSeg’s probability maps are sharply bimodal: background voxels are driven to near-zero probability and foreground voxels to near-one, with a narrow transition band between them (Fig. 2). The resulting segmentation is therefore largely insensitive to the exact threshold. The specialist models are less well calibrated by comparison: they assign appreciable probability to background and produce blurry, gradual foreground-background transitions, so their binarized output depends more strongly on the chosen threshold. This sharper separation of foreground from background indicates that SubCellSeg is more certain about what is signal and what is not. Across the 13 structures we tested, the 0.5 threshold yielded near-optimal F1 across the validation set; structure-, study-, and application-specific thresholds can yield additional gains and can be calibrated using the utility function provided in the `allencell-segmenter-ml` package.

#### Evaluation metrics

Segmentation performance was reported using the Dice similarity coefficient (DSC): Dice = 2*|P ∩ G|/*(*|P |* + *|G|*), where *P* is the set of predicted foreground voxels and *G* is the set of reference mask foreground voxels. Dice is reported as the mean over test images *±* standard deviation. We used Dice because it measures foreground overlap only and is therefore not dominated by background, which occupies most of each volume for sparse structures.

#### Reproducibility

The SubCellSeg model, checkpoint, training pipeline, and inference tooling are distributed as part of the allencell-segmenter-ml package (https://github.com/AllenCell/allencell-segmenter-ml), and the benchmarking, evaluation, and figure-generation code for this study, including the specialist and cross-evaluation experiments, are available at https://github.com/AllenCell/SubCellSeg.

#### Statistics

Per-structure Dice scores are reported as mean *±* standard deviation across the held-out test set (57 images per structure, 741 images in total). Bars in Fig. 1**d** show the per-structure mean, and error bars give *±*1 standard deviation. Because every test image is segmented by every method, between-method comparisons are paired at the image level. Significance was assessed with two-sided paired Wilcoxon signed-rank tests on per-image Dice, comparing SubCellSeg against each competing method (the structure-specific specialist models and the published baselines) separately for each structure. All such comparisons were significant (*p <* 10*^−^*^7^ for every structure and method).

## Reporting Summary

Further information on research design is available from the corresponding author on request.

## Data and Code Availability

Training images and reference segmentations are part of the WTC-11 hiPSC Single-Cell Image Dataset v1 [2] (https://open.quiltdata.com/b/allencell). The cell lines are available through the Allen Cell Collection (https://www.allencell.org/). SubCellSeg model code and pretrained checkpoints for all four backbone architectures are available at https://github.com/AllenCell/allencell-segmenter-ml, distributed under the Allen Institute Software License. The plugin can be installed via `pip install allencell-segmenter-ml`. A command-line interface supports batch inference on local image collections without requiring the napari GUI. The nucleoli DFC drug-perturbation images are part of the dataset published with [3]. The remaining drug-perturbation images, the cross-microscope and cardiomyocyte test images, and all model predictions will be made public upon publication; until then, they are available from the corresponding author upon reasonable request.

## Acknowledgments

We thank members of the Allen Institute for Cell Science for generating and curating the training data, developing the classical segmentation workflows that served as training labels, and providing feedback on the model and interface.

