## Supplementary file for "SubCellSeg: A generalist model for subcellular structure segmentation in 3D fluorescence microscopy"

### Supplementary Information for
### SubCellSeg: A human annotation-free generalist model for subcellular structure segmentation in 3D fluorescence microscopy

#### Contents

**Extended Data Figures** 2

**Supplementary Notes** 8

Note 1 — Test-set curation . . . . . . . . . . 8

Note 2 — Backbone configuration rationale . . . . . . . . . . 8

**Supplementary Tables** 9

### Extended Data Figures

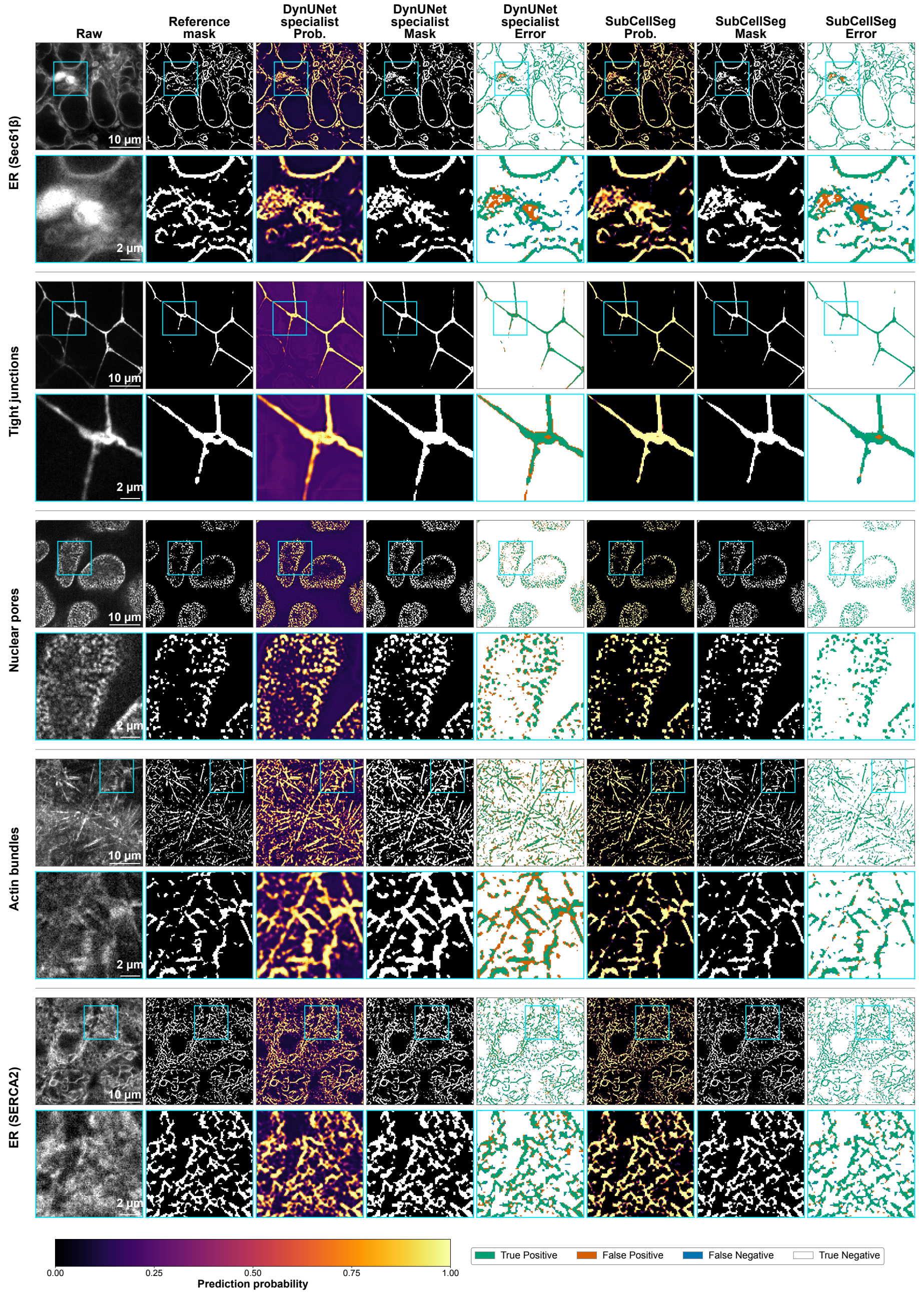


**Extended Data Figure 1. Qualitative segmentation results for five further trained structures (part 1 of 2).** Companion to Fig. 2 of the main text. Structures shown here: ER via Sec61$\beta$, tight junctions via ZO-1, nuclear pores via NUP153, actin bundles via $\alpha$-actinin-1 and ER via SERCA2. Layout as in Fig. 2: for each structure, the full field of view (top row) and a 3.2$\times$ magnification of the cyan boxed region (bottom row), with columns showing the raw image, the reference mask, the DynUNet specialist probability map, binary mask and error map, and the SubCellSeg probability map, binary mask and error map. Probability maps share the colour scale at bottom (prediction probability 0–1). Error maps use green, true positive; orange, false positive; blue, false negative; white, true negative. Scale bars, 10 $\mu$m (full view) and 2 $\mu$m (magnified view).

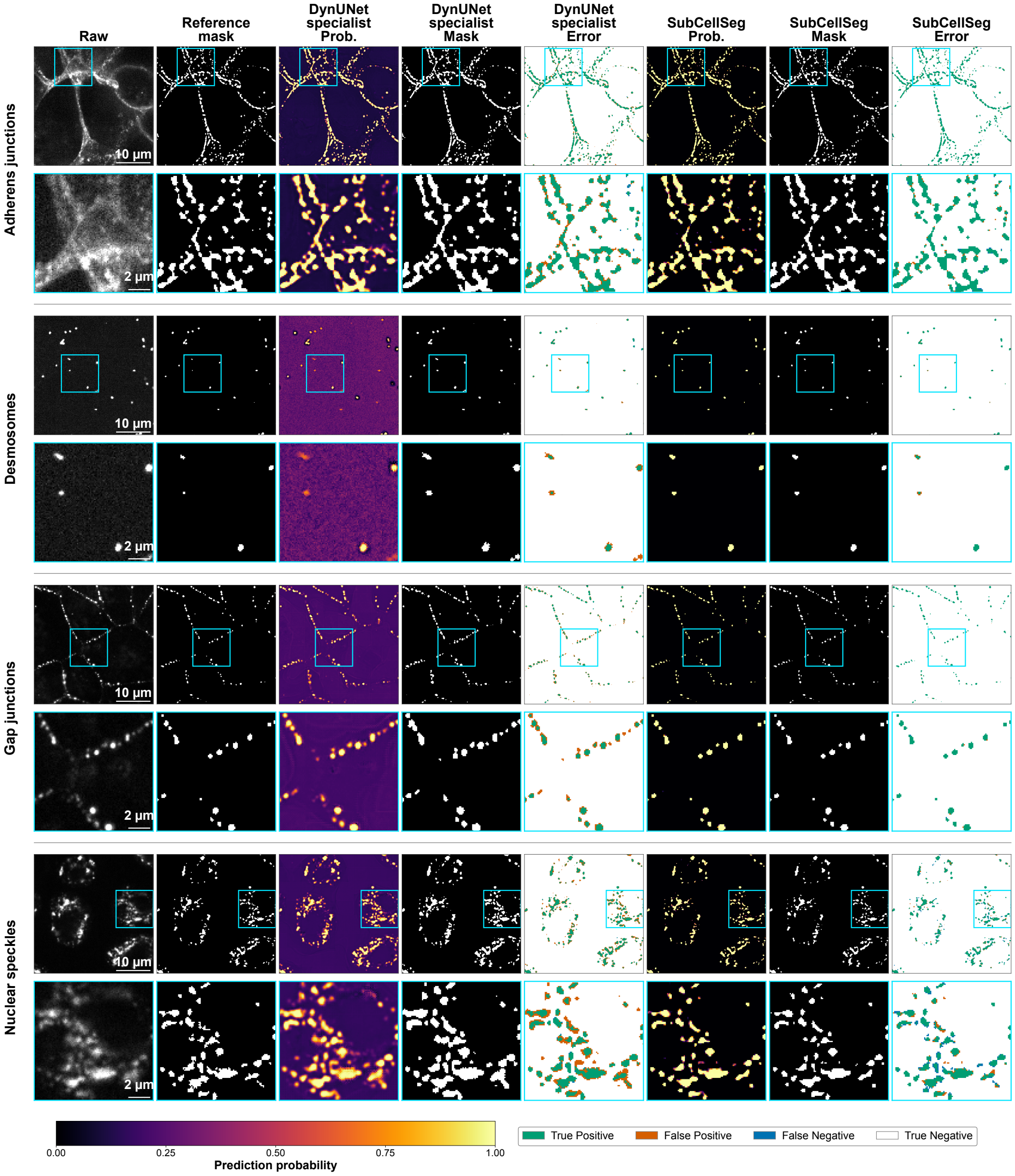


**Extended Data Figure 1. Qualitative segmentation results for the remaining four trained structures (part 2 of 2).** Adherens junctions via $\beta$-catenin, desmosomes via desmoplakin, gap junctions via connexin-43 and nuclear speckles via SON. Layout, columns, cyan magnification box and error-map colour code as in Fig. 2 and Extended Data Fig. 1 above; together the three figures cover all 13 training structures. Probability maps share the colour scale at bottom (prediction probability 0–1). Scale bars, 10 $\mu$m (full view) and 2 $\mu$m (magnified view).

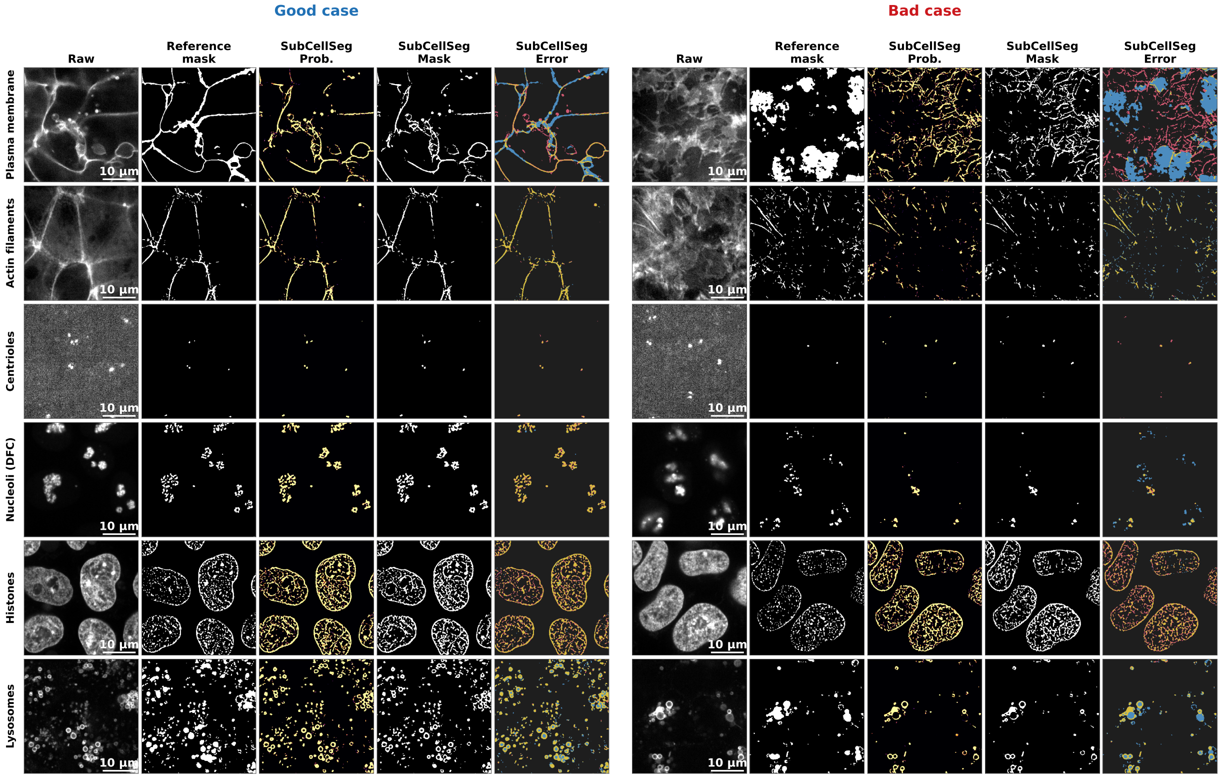


**Extended Data Figure 2. Zero-shot performance on 12 additional unseen subcellular structures (part 1 of 2).** SubCellSeg applied without any fine-tuning to 12 structures from the Allen Cell Collection that were absent from its training set, imaged on the same spinning-disk system at $100\times$. For each structure, a representative "good case" (left five columns) and "bad case" (right five columns) are shown: raw image, reference mask, SubCellSeg probability map, SubCellSeg binary mask, and SubCellSeg error map. No specialist model is shown, as none exists for these structures. Error maps use yellow: true positive; red: false positive; blue: false negative; black: true negative). Good and bad slices were selected automatically as the highest- and lowest-Dice foreground-rich $z$-slices of the same volume. This part shows plasma membrane via CAAX, actin filaments via $\beta$-actin, centrioles via centrin-2, nucleoli (DFC) via fibrillarin, histones via histone H2B and lysosomes via LAMP-1. Scale bars, 10 $\mu$m.

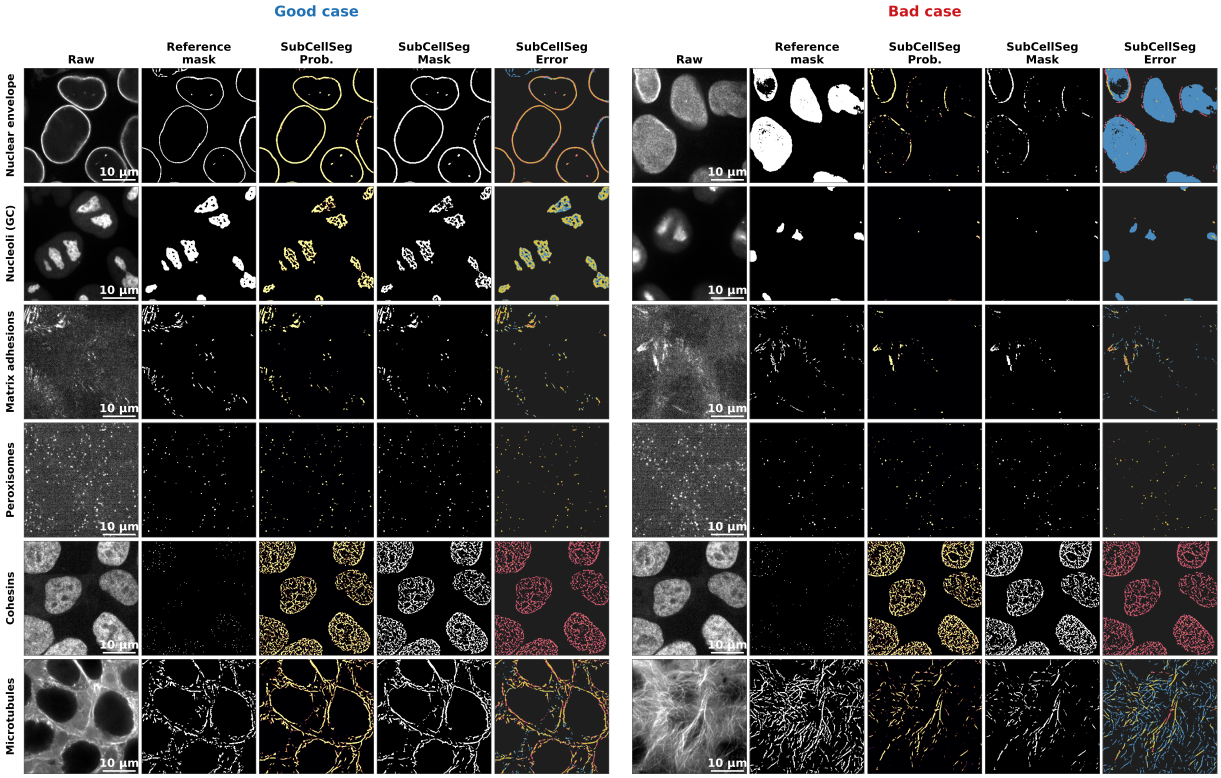


**Extended Data Figure 2. Zero-shot performance on 12 additional unseen subcellular structures (part 2 of 2).** Continued from the previous page. This part shows nuclear envelope via lamin B1, nucleoli (GC) via nucleophosmin, matrix adhesions via paxillin, peroxisomes via PMP34, cohesins via SMC1A and microtubules via $\alpha$-tubulin. Columns, colour code and slice selection as in part 1. Scale bars, 10 $\mu$m.

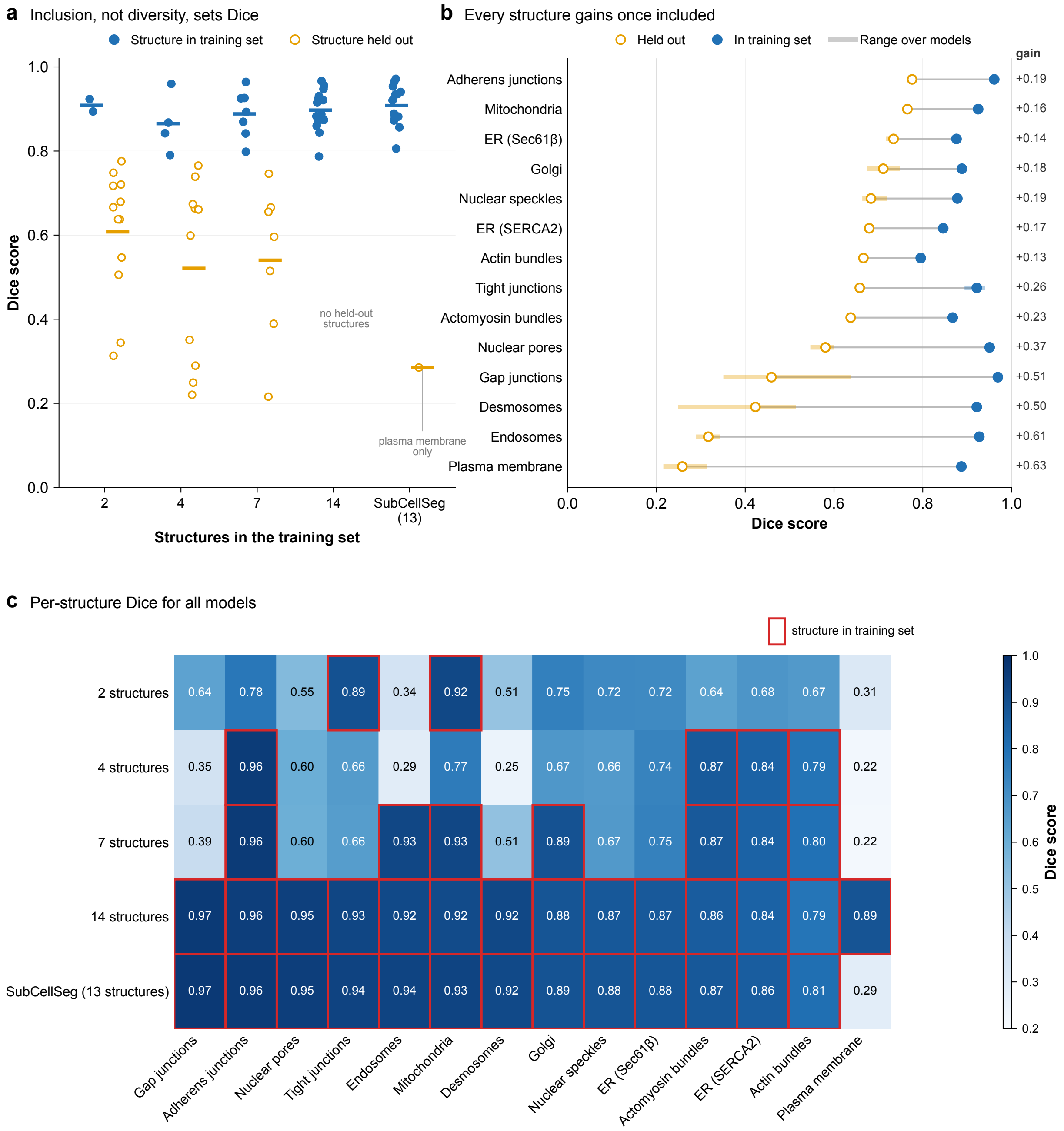


**Extended Data Figure 3. Segmentation accuracy depends on whether a structure was in the training set, not on how many structures were.** Models were trained from scratch on 2 (mitochondria via TOM20, tight junctions via ZO-1), 4, 7 or 14 structures and compared with SubCellSeg (13 structures). The fourteenth structure is the plasma membrane via CAAX (AAVS1 safe-harbour line), which SubCellSeg never saw. All values are mean Dice over the 57-image filtered test cohort of each structure. (**a**) Per-model Dice for structures in that model's training set (filled blue) and held out from it (open orange); dots are structures, bars the mean over structures. In-training Dice is 0.86–0.91 regardless of training-set size, and held-out Dice is 0.52–0.61 and non-monotonic: the 2-structure model generalizes better than the 7-structure model (0.61 versus 0.54). SubCellSeg's only held-out structure is the plasma membrane (Dice 0.29). (**b**) Per-structure Dice when held out (open orange) versus when included (filled blue); markers are means over the models in each group, bands the range, gain printed at right. Every structure improves when included (+0.13 to +0.63); sparse punctate structures (endosomes, desmosomes, plasma membrane, gap junctions) transfer worst when unseen. (**c**) Full matrix of mean Dice; red outlines mark pairs where the structure was in the training set. Adding the plasma membrane (14-structure model) raises its own Dice from $\leq 0.31$ to 0.89 and lowers the 13 shared structures by at most 0.02 (mean 0.90 versus 0.91 for SubCellSeg). Models should therefore be compared per structure on the 13 shared targets: the 14-structure average favours the 14-structure model only because of the plasma-membrane term. Means in **a** and **b** are over structures and rest on different numbers of structures per model.

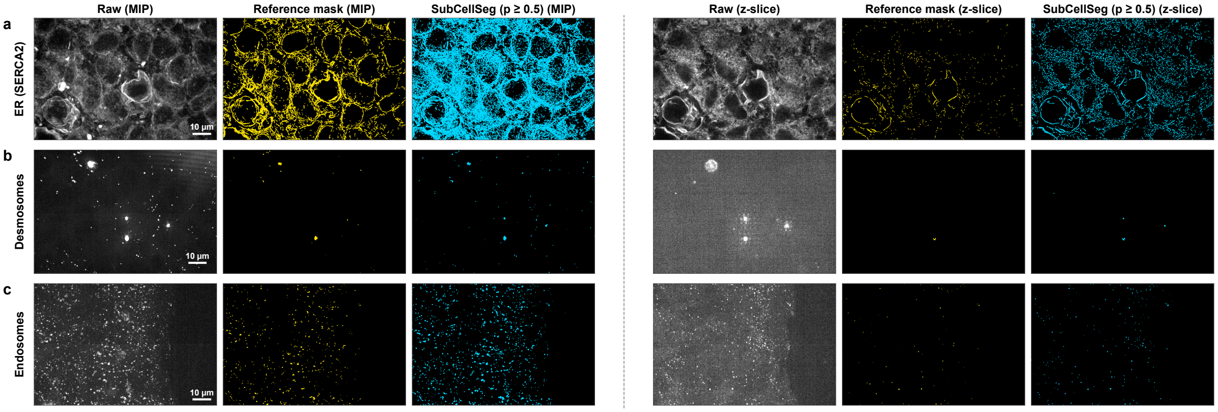


**Extended Data Figure 4. Examples of incomplete classical reference masks among the excluded test images.** Three fields of view from the test images excluded from the benchmark (the 10 lowest-scoring images per structure; Methods), each shown as raw image | reference mask (classical Allen Cell and Structure Segmenter workflow) | SubCellSeg prediction (probability $\geq 0.5$), as a maximum-intensity $z$-projection (left block) and at a single representative $z$-slice (right block). Reference and prediction masks are overlaid in gold and cyan; scale bar, $10\,\mu$m. (**a**) ER via SERCA2 (image ATP2A2_146596): the reference mask covers only part of the reticular network visible in the raw image, which SubCellSeg segments more fully (Dice $= 0.36$, recall $= 1.00$, precision $= 0.22$). (**b**) Desmosomes via desmoplakin (image DSP_4596), the structure most affected and the one for which biologists confirmed the limitation of the reference workflow: the reference mask marks only a few of the desmosome puncta, while SubCellSeg detects the broader punctate pattern present in the raw image (Dice $= 0.52$, recall $= 0.93$, precision $= 0.36$). (**c**) Endosomes via RAB5A (image RAB5A_149458): the reference mask labels only a fraction of the RAB5A puncta visible in the raw image, whereas SubCellSeg recovers nearly all of them (Dice $= 0.43$, recall $= 1.00$, precision $= 0.27$). In each case, the low Dice arises mainly from structure that is visible in the raw image but absent from the reference mask: recall against the reference is close to 1, while precision is low. The excluded images, their reference masks, and all predictions are publicly available (Data and Code Availability).

### Supplementary Notes

#### Supplementary Note 1 | Test-set curation and outlier identification

The Methods describe a reduction of the held-out test set from 67 to 57 z-stacks per structure. Because the reduction used model output as part of the selection signal, this note states the procedure in full and reports performance both before and after the reduction, so that the reported improvement can be checked against the unreduced test set.

**Motivation.** During evaluation, a small number of test images showed unusually low segmentation accuracy. In these images, the reference mask, generated by the classical Allen Cell and Structure Segmenter workflow, captured only part of the structure visible in the raw image. The problem was most pronounced for desmosomes via desmoplakin, where biologists confirmed the limitation of the reference workflow, and similar cases occurred for other structures (Extended Data Fig. 4).

**Selection procedure.** For each of the 13 training structures, test images were ranked by Dice score computed from the SubCellSeg-DynUNet predictions against the reference masks. The 10 lowest-scoring images per structure were flagged for removal. Those same sample IDs were then excluded from all evaluation types (individual specialist, SubCellSeg with every backbone, cross-evaluation) so that every method is assessed on an identical 57-image subset. The same number of images (10) was removed from every structure to keep the evaluation balanced, giving 57 z-stacks per structure (741 images across 13 structures).

**Agreement across backbones.** The images identified were largely independent of which model was used to rank them. When the same bottom-10 procedure was run independently with each of the four SubCellSeg backbones (DynUNet, AttentionUNet, SwinUNETR, UNet), the pairwise overlap between any two backbones' exclusion sets ranged from 85% to 92% (mean $\sim$90%); 108 of the 130 DynUNet flagged samples (83%) were also flagged by all three remaining backbones. Because a poor reference mask degrades every model that is scored against it, this consistency indicates the flagged images reflect annotation quality rather than a failure mode specific to one architecture.

**Effect on reported performance.** The table below is the substantive check: whether the SubCellSeg-over-specialist improvement depends on the curation. The SubCellSeg advantage is preserved on the unreduced 67-image test set (+0.124) and is comparable to the curated result (+0.136). Both models benefit from curation (each gains $\sim$0.01 in absolute Dice), confirming that the removed images penalize all methods equally and the relative ranking is not an artifact of filtering.

**Backbone configuration rationale.** The backbone comparison is intended as a practical survey of architectures a user of SubCellSeg's CytoDL-based training pipeline would encounter, not as a parameter-controlled ablation. Each backbone was configured at its MONAI recommended or default setting:

- **DynUNet** computes encoder channel widths automatically from the input strides ($\texttt{filters}[i] = \min(2^{5+i}, 320)$), yielding a five-stage encoder with channels $[32, 64, 128, 256, 320]$ and residual blocks enabled. This resolution-adaptive sizing is a defining feature of the architecture [1] and was retained as-is.
- **SwinUNETR** was run at $\texttt{feature\_size} = 48$, double MONAI's default of 24, following the setting recommended in [2].
- **AttentionUNet** and **UNet** use $\texttt{channels} = [16, 32, 64, 128, 256]$, the configuration in MONAI's 3D segmentation tutorials and the CytoDL starter template.

Because these are default rather than size-matched configurations, the parameter counts differ substantially (4.8 M–63.1 M). The comparison therefore conflates architectural inductive biases with model capacity: scaling AttentionUNet or UNet to DynUNet-level channel widths (e.g., $[32, 64, 128, 256, 320]$) might narrow the Dice gap, though SwinUNETR's under-performance at nearly double DynUNet's parameter count suggests that capacity alone does not determine performance on this task. We report the comparison at default settings because it reflects the out-of-the-box experience for practitioners adopting the pipeline, and because the paper's central claim: *SubCellSeg versus specialist models*, is independent of backbone choice (all four backbones outperform the specialist baseline).

#### Supplementary Tables

**Supplementary Table 1. Performance before and after test-set curation.** Mean Dice across the 13 trained structures, evaluated on the full test set (67 z-stacks per structure) and on the curated test set (57 per structure).

| Method | Full set (67/structure) | Curated (57/structure) |
|---|---|---|
| SubCellSeg (DynUNet) | 0.888 | 0.909 |
| DynUNet specialists | 0.764 | 0.773 |
| **Difference** | **+0.124** | **+0.136** |

**Supplementary Table 2. Per-structure Dice for all evaluated methods.** Mean Dice on the held-out test set (57 images per structure). Column means appear in Table 2 of the main text. Gene names in parentheses identify the tagged protein; see Supplementary Table 5.

| Structure | SubCellSeg | nnU-Net | Special. | Otsu | Nellie | CellPose | StarDist | MicroSAM | SAM |
|---|---|---|---|---|---|---|---|---|---|
| Endosomes (RAB5A) | 0.936 | 0.837 | 0.667 | 0.010 | 0.526 | 0.214 | 0.195 | 0.000 | 0.004 |
| Gap junctions (GJA1) | 0.972 | 0.900 | 0.726 | 0.522 | 0.732 | 0.179 | 0.118 | 0.003 | 0.004 |
| Desmosomes (DSP) | 0.926 | 0.874 | 0.689 | 0.183 | 0.752 | 0.015 | 0.166 | 0.000 | 0.001 |
| Nuclear pores (NUP153) | 0.954 | 0.843 | 0.785 | 0.276 | 0.644 | 0.127 | 0.080 | 0.111 | 0.024 |
| Tight junctions (TJP1) | 0.940 | 0.877 | 0.784 | 0.790 | 0.749 | 0.461 | 0.278 | 0.027 | 0.003 |
| Actomyosin (MYH10) | 0.873 | 0.769 | 0.723 | 0.263 | 0.369 | 0.141 | 0.246 | 0.061 | 0.024 |
| Actin bundles (ACTN1) | 0.807 | 0.750 | 0.675 | 0.101 | 0.630 | 0.074 | 0.199 | 0.055 | 0.020 |
| Nuclear speckles (SON) | 0.882 | 0.828 | 0.781 | 0.663 | 0.726 | 0.358 | 0.414 | 0.004 | 0.009 |
| Golgi (ST6GAL1) | 0.889 | 0.859 | 0.798 | 0.078 | 0.609 | 0.602 | 0.522 | 0.000 | 0.014 |
| Adherens junctions (CTNNB1) | 0.965 | 0.831 | 0.883 | 0.332 | 0.495 | 0.200 | 0.176 | 0.053 | 0.013 |
| ER (ATP2A2) | 0.857 | 0.821 | 0.794 | 0.291 | 0.481 | 0.098 | 0.235 | 0.022 | 0.068 |
| ER (SEC61B) | 0.881 | 0.849 | 0.832 | 0.348 | 0.436 | 0.094 | 0.227 | 0.079 | 0.061 |
| Mitochondria (TOMM20) | 0.935 | 0.892 | 0.915 | 0.685 | 0.651 | 0.707 | 0.351 | 0.000 | 0.077 |
| **Mean** | **0.909** | 0.841 | 0.773 | 0.350 | 0.600 | 0.251 | 0.249 | 0.032 | 0.025 |

**Supplementary Table 3. Model size and performance by backbone.** All models trained under the identical protocol described in Methods (1000 epochs, NVIDIA A100 80 GB, PyTorch 2.4.0). Mean Dice is computed on the curated test set (57 images × 13 structures). Each backbone was instantiated via MONAI (v1.3) with its recommended configuration (see text below). Parameter counts were measured from the released checkpoints.

| Backbone | Parameters | Mean Dice | Configuration source |
|---|---|---|---|
| DynUNet | 33.3 M | **0.909** | MONAI auto-filters $[32, 64, 128, 256, 320]$ |
| SwinUNETR | 63.1 M | 0.904 | `feature_size` $= 48$ |
| AttentionUNet | 5.9 M | 0.892 | `channels` $= [16, 32, 64, 128, 256]$ |
| UNet | 4.8 M | 0.870 | `channels` $= [16, 32, 64, 128, 256]$ |

**Supplementary Table 4. Structure-specific and promptable baselines.** Each tool was evaluated zero-shot on the trained structures matching its design (57 held-out images per structure). The score is voxel-wise Dice, except for Spotiflow rows. Spotiflow outputs spot coordinates rather than masks, so those rows use object-level detection F1 for both methods. Each mask is reduced to the centroids of its connected components and matched one-to-one to reference mask centroids within 1 $\mu$m (Methods). This F1 counts objects, not voxels, so it is not the same quantity as Dice, which equals voxel-wise F1. SubCellSeg's detection F1 (DSP 0.944, RAB5A 0.931) therefore differs from its Dice on the same images (0.926, 0.936). nnInteractive "oracle" prompts are derived from the reference masks (bounding box and corrective clicks; Methods). Values are mean ± s.d. In every row, SubCellSeg (DynUNet) scored higher on all 57 images ($p < 10^{-10}$, two-sided paired Wilcoxon signed-rank test). The Gene column gives the tagged gene for each structure; see Supplementary Table 5.

| Method | Design target | Gene | Metric | Method score | SubCellSeg |
|---|---|---|---|---|---|
| ERnet | ER (2D SIM) | ATP2A2 | Dice | 0.089 ± 0.016 | 0.857 ± 0.015 |
| | | SEC61B | Dice | 0.134 ± 0.025 | 0.881 ± 0.018 |
| PlantSeg (generic confocal) | Cell boundaries | CTNNB1 | Dice | 0.213 ± 0.030 | 0.965 ± 0.003 |
| Spotiflow (smFISH 3D) | Spots | DSP | F1 | 0.354 ± 0.064 | 0.944 ± 0.037 |
| | | RAB5A | F1 | 0.552 ± 0.046 | 0.931 ± 0.025 |
| Spotiflow (synthetic 3D) | Spots | DSP | F1 | 0.033 ± 0.014 | 0.944 ± 0.037 |
| | | RAB5A | F1 | 0.201 ± 0.044 | 0.931 ± 0.025 |
| nnInteractive, FOV box | Promptable 3D | ACTN1 | Dice | 0.028 ± 0.012 | 0.807 ± 0.034 |
| | | ATP2A2 | Dice | 0.098 ± 0.031 | 0.857 ± 0.015 |
| | | CTNNB1 | Dice | 0.032 ± 0.006 | 0.965 ± 0.003 |
| | | DSP | Dice | 0.001 ± 0.000 | 0.926 ± 0.041 |
| | | GJA1 | Dice | 0.004 ± 0.000 | 0.972 ± 0.003 |
| | | MYH10 | Dice | 0.028 ± 0.008 | 0.873 ± 0.014 |
| | | NUP153 | Dice | 0.021 ± 0.006 | 0.954 ± 0.008 |
| | | RAB5A | Dice | 0.004 ± 0.002 | 0.936 ± 0.026 |
| | | SEC61B | Dice | 0.115 ± 0.026 | 0.881 ± 0.018 |
| | | SON | Dice | 0.009 ± 0.001 | 0.882 ± 0.006 |
| | | ST6GAL1 | Dice | 0.014 ± 0.004 | 0.889 ± 0.016 |
| | | TJP1 | Dice | 0.007 ± 0.001 | 0.940 ± 0.008 |
| | | TOMM20 | Dice | 0.079 ± 0.014 | 0.935 ± 0.017 |
| | | *Mean (13)* | Dice | 0.034 | 0.909 |
| nnInteractive, oracle box | Promptable 3D | *Mean (13)* | Dice | 0.035 | 0.909 |
| nnInteractive, oracle box + 5 clicks | Promptable 3D | *Mean (13)* | Dice | 0.036 | 0.909 |

**Supplementary Table 5. Cell lines.** All lines are WTC-11 hiPSCs with an endogenously tagged marker protein (references in Methods). All lines except AICS-0007 are available through the Allen Cell Collection. "Comparisons" means the specialist and state-of-the-art comparisons; "unseen structures" means zero-shot evaluation on 12 untrained structures (Extended Data Fig. 2); "zero-shot imaging" means other microscopes, magnifications or cell types (Fig. 6).

| Cell line | Clone | Tagged allele | Tagged protein | Structure | Used in |
|---|---|---|---|---|---|
| AICS-0007 | cl.79 | ACTN1-mEGFP | $\alpha$-actinin-1 | actin bundles | training; comparisons |
| AICS-0010 | cl.55 | mEGFP-SEC61B | Sec61$\beta$ | ER (SEC61$\beta$) | training; comparisons |
| AICS-0011 | cl.27 | TOMM20-mEGFP | TOM20 | mitochondria | training; comparisons; perturbation |
| AICS-0017 | cl.65 | DSP-mEGFP | desmoplakin | desmosomes | training; comparisons |
| AICS-0023 | cl.20 | mEGFP-TJP1 | ZO-1 | tight junctions | training; comparisons |
| AICS-0024 | cl.80 | mEGFP-MYH10 | non-muscle myosin IIB | actomyosin bundles | training; comparisons |
| AICS-0025 | cl.1; cl.44 | ST6GAL1-mEGFP | sialyltransferase 1 | Golgi | training; comparisons; perturbation |
| AICS-0040 | cl.13; cl.35 | mEGFP-RAB5A | RAB5A | endosomes | training; comparisons |
| AICS-0046 | cl.51 | mEGFP-ATP2A2 | SERCA2 | ER (SERCA2) | training; comparisons |
| AICS-0053 | cl.16 | GJA1-mEGFP | connexin-43 | gap junctions | training; comparisons |
| AICS-0058 | cl.67 | mEGFP-CTNNB1 | $\beta$-catenin | adherens junctions | training; comparisons |
| AICS-0069 | cl.88 | mEGFP-NUP153 | NUP153 | nuclear pores | training; comparisons; perturbation |
| AICS-0094 | cl.24 | mEGFP-SON | SON | nuclear speckles | training; comparisons |
| AICS-0005 | cl.50 | PXN-EGFP | paxillin | matrix adhesions | unseen structures |
| AICS-0012 | cl.105 | mEGFP-TUBA1B | $\alpha$-tubulin | microtubules | unseen structures; perturbation; zero-shot imaging |
| AICS-0013 | cl.210 | mEGFP-LMNB1 | lamin B1 | nuclear envelope | unseen structures; fine-tuning |
| AICS-0014 | cl.6 | FBL-mEGFP | fibrillarin | nucleolus (DFC) | unseen structures; perturbation |
| AICS-0016 | cl.184 | mEGFP-ACTB | $\beta$-actin | actin filaments | unseen structures |
| AICS-0022 | cl.37 | LAMP1-mEGFP | LAMP-1 | lysosomes | unseen structures; fine-tuning |
| AICS-0032 | cl.19 | mTagRFP-T-CETN2 | centrin-2 | centrioles | unseen structures |
| AICS-0033 | cl.115 | SLC25A17-mEGFP | PMP34 | peroxisomes | unseen structures |
| AICS-0054 | cl.91 | mTagRFP-T-CAAX | CAAX (K-Ras) | plasma membrane | unseen structures |
| AICS-0057 | cl.50 | NPM1-mEGFP | nucleophosmin | nucleolus (GC) | unseen structures; zero-shot imaging |
| AICS-0061 | cl.36 | HIST1H2BJ-mEGFP | histone H2B type 1-J | histones | unseen structures |
| AICS-0068 | cl.9 | SMC1A-mEGFP | SMC1A | cohesins | unseen structures |
| AICS-0088 | cl.83 | mEGFP-PCNA | PCNA | DNA replication foci | zero-shot imaging |
| AICS-0095 | cl.18 | mEGFP-EZH2 | EZH2 | PRC2 | zero-shot imaging |
| AICS-0048 | cl.39 | TTN-mEGFP | titin | sarcomeres (M-line), cardiomyocytes | zero-shot imaging |

**Supplementary Table 6. Acquisition parameters of the zero-shot images in Fig. 6.** Microscope system and configuration details for images in each panel. Laser information is provided for only the channel(s) displayed in the panel corresponding to 488 nm unless otherwise noted; additional lasers may have been used during image acquisition for channels omitted from this study (see Methods).

| Panel | Protein | Line | System | Objective | Exposure / pixel dwell | Laser power (%) | Voxel size |
|---|---|---|---|---|---|---|---|
| 6b top | PCNA | AICS-0088 | SDCM-1 | 100×/1.25 W | 200 ms | 10 | 0.108×0.108×0.290 |
| 6b bottom | EZH2 | AICS-0095 | SDCM-1 | 100×/1.25 W | 200 ms | 41 | 0.108×0.108×0.290 |
| 6c top | nucleophosmin | AICS-0057 | SDCM-1 | 20×/0.8 | 200 ms | 20 | 0.271×0.271×0.530 |
| 6c middle | nucleophosmin | AICS-0057 | LSCM | 40×/1.2 W | 0.55 µs with 2x line averaging | 1 | 0.083×0.083×0.208 |
| 6c bottom | nucleophosmin | AICS-0057 | SDCM-1 | 100×/1.25 W | 200 ms | 10 | 0.108×0.108×0.290 |
| 6d | $\alpha$-tubulin | AICS-0012 | LSCM | 40×/1.2 W | 0.55 µs with 2x line averaging | 5 | 0.083×0.083×0.208 |
| 6e(i) | titin (live) | AICS-0048 | SDCM-1 | 40×/1.2 W | 100 ms | 89.8 | 0.135×0.135×0.290 |
| 6e(ii) | titin (Z-line Ab) | AICS-0048 | SDCM-2 | 63×/1.2 W, 0.83× tube lens | 100 ms (for each of 488/561/640) | 488/561/640 at 50, 10, and 30 | 0.124×0.124×0.33 |
| 6e(iii) | titin (M-line Ab) | AICS-0048 | SDCM-2 | 63×/1.2 W, 0.83× tube lens | 100 ms (for each of 488/561/640) | 488/561/640 at 50, 10, and 30 | 0.124×0.124×0.33 |

**Supplementary Table 7. Structure-diversity ablation on the 25-structure held-out test set.** Mean Dice of models trained from scratch on 2, 4, 7 or 14 structures and of SubCellSeg (13 structures), evaluated on two images per structure (50 images; separate from the 57-image cohorts of Extended Data Fig. 3). Bold marks structures that were in that model's training set. The 2-structure model was trained on mitochondria and tight junctions; the 4-structure model on actin bundles, ER (SERCA2), adherens junctions and actomyosin bundles; the 7-structure model on those four plus endosomes, Golgi and mitochondria.

| Structure | 2 structures | 4 structures | 7 structures | 14 structures | SubCellSeg (13) |
|---|---|---|---|---|---|
| *SubCellSeg training structures* | | | | | |
| Gap junctions | 0.65 | 0.36 | 0.40 | **0.97** | **0.97** |
| Adherens junctions | 0.77 | **0.95** | **0.95** | **0.91** | **0.81** |
| Nuclear pores | 0.66 | 0.68 | 0.68 | **0.96** | **0.96** |
| Tight junctions | **0.90** | 0.66 | 0.67 | **0.94** | **0.95** |
| Endosomes | 0.36 | 0.32 | **0.95** | **0.95** | **0.96** |
| Mitochondria | **0.90** | 0.81 | **0.92** | **0.92** | **0.91** |
| Desmosomes | 0.54 | 0.29 | 0.48 | **0.96** | **0.97** |
| Golgi | 0.65 | 0.60 | **0.84** | **0.83** | **0.82** |
| Nuclear speckles | 0.71 | 0.66 | 0.66 | **0.87** | **0.88** |
| ER (Sec61$\beta$) | 0.71 | 0.76 | 0.76 | **0.87** | **0.88** |
| Actomyosin bundles | 0.61 | **0.87** | **0.87** | **0.86** | **0.87** |
| ER (SERCA2) | 0.69 | **0.86** | **0.85** | **0.86** | **0.87** |
| Actin bundles | 0.70 | **0.72** | **0.70** | **0.73** | **0.69** |
| *Added in the 14-structure model only* | | | | | |
| Plasma membrane (CAAX) | 0.27 | 0.18 | 0.17 | **0.86** | 0.23 |
| *Unseen by every model* | | | | | |
| Actin filaments | 0.55 | 0.79 | 0.78 | 0.20 | 0.78 |
| Centrioles | 0.30 | 0.11 | 0.28 | 0.59 | 0.59 |
| Nucleoli (DFC) | 0.72 | 0.77 | 0.77 | 0.72 | 0.77 |
| Histones | 0.59 | 0.75 | 0.75 | 0.66 | 0.67 |
| Lysosomes | 0.75 | 0.74 | 0.76 | 0.75 | 0.73 |
| Nuclear envelope | 0.32 | 0.29 | 0.28 | 0.61 | 0.31 |
| Nucleoli (GC) | 0.78 | 0.55 | 0.55 | 0.34 | 0.58 |
| Matrix adhesions | 0.66 | 0.32 | 0.54 | 0.59 | 0.55 |
| Peroxisomes | 0.29 | 0.20 | 0.88 | 0.83 | 0.87 |
| Cohesins | 0.04 | 0.05 | 0.05 | 0.03 | 0.04 |
| Microtubules | 0.49 | 0.24 | 0.26 | 0.51 | 0.50 |
| *Mean over the 11 unseen structures* | *0.50* | *0.44* | *0.54* | *0.53* | *0.58* |
